# Assessment of ethanol and nicotine interactions using a reinforcer demand modeling with grouped and individual levels of analyses in a long-access self-administration model using male rats

**DOI:** 10.1101/2022.10.17.512519

**Authors:** Christopher L. Robisona, Nicole Covab, Victoria Madoreb, Tyler Allenb, Scott Barrettc, Sergios Charntikov

## Abstract

Previous reports have indicated the reciprocal effects of nicotine and ethanol on their rewarding and reinforcing properties, but studies using methodological approaches resembling substance use in vulnerable populations are lacking. In our study, rats first self-administered ethanol, and their sensitivity to ethanol’s reinforcing effects was assessed using a reinforcer demand modeling approach. Subsequently, rats were equipped with intravenous catheters to self-administer nicotine, and their sensitivity to nicotine’s reinforcing effects was evaluated using the same approach. In the final phase, rats were allowed to self-administer ethanol and nicotine concurrently, investigating the influence of one substance on the rate of responding for the other substance. Group analyses revealed notable differences in demand among sucrose, sweetened ethanol, and ethanol-alone, with sucrose demonstrating the highest demand and ethanol-alone exhibiting greater sensitivity to changes in cost. At the individual level, our study finds significant correlations between rats’ demand for sucrose and sweetened ethanol, suggesting parallel efforts for both substances. Our individual data also suggest interconnections in the elasticity of demand for sweetened ethanol and ethanol-alone, as well as a potential relationship in price response patterns between ethanol and nicotine. Furthermore, concurrent self-administration of ethanol and nicotine at the group level displayed reciprocal effects, with reduced responding for nicotine in the presence of ethanol and increased responding for ethanol in the presence of nicotine. This study provides valuable insights into modeling the co-use of ethanol and nicotine and assessing their interaction effects using reinforcer demand modeling and concurrent self-administration or noncontingent administration tests. These findings contribute to our understanding of the complex interplay between ethanol and nicotine and have implications for elucidating the underlying mechanisms involved in polydrug use.

## 1. Introduction

Tobacco and alcohol consumption rank among the top contributors to preventable fatalities globally. Collectively, these substances are attributable to approximately 11 million avoidable deaths annually - 8 million and 3 million for tobacco and alcohol, respectively (Murray et al., 2020; WHO, 2018). Moreover, the co-occurrence of these substance uses is alarmingly prevalent (EMCDDA, 2009; Kohut, 2017). For instance, it has been found that nearly 90 % of patients diagnosed with alcohol use disorder also report regular tobacco use (Burling and Ziff, 1988; DiFranza and Guerrera, 1990; Toneatto et al., 1995). Recent studies indicate that the combined use of tobacco and alcohol inflicts more harm on users than the individual use of either substance (Frie et al., 2021; Hurt et al., 1996; Kohut, 2017; Mello et al., 1980; Mintz et al., 1985). Moreover, it has been suggested that polydrug users might face heightened difficulties in obtaining effective treatment for one of the substances they use (Kozlowski et al., 1989; Stuyt, 1997). Presently, the treatments available for substance use disorders demonstrate limited effectiveness, and they fall short in providing personalized approaches that take into account an individual’s specific history of polydrug use. Although it is crucial to evaluate group effects in research studies for broader understanding, relying solely on average markers of performance may not yield accurate information about the diverse mechanisms at play at an individual level. Therefore, there is an exigent need to deepen our understanding of the interactions between nicotine and alcohol use, with a particular emphasis on individual differences.

The interactions between nicotine and ethanol have been a topic of extensive exploration in both clinical and preclinical fields. These studies employ diverse methodological approaches, yielding somewhat inconsistent findings regarding the effects of nicotine on alcohol reinforcement. Clinical research indicates that nicotine escalates alcohol consumption and intensifies the effort participants are willing to exert for alcohol reinforcement (Barrett et al., 2006; Dermody et al., 2016). However, preclinical studies present varied outcomes regarding nicotine’s impact on alcohol reinforcement. These discrepancies may arise from the diverse methodologies used to gather relevant data. For instance, some studies depend on noncontingent modes of ethanol delivery, such as bottle or vapor (Chandler et al., 2020; Lallemand et al., 2007; Marshall et al., 2003; Potthoff et al., 1983; Smith et al., 1999), while others employ contingent operant protocols for self-administration of ethanol (Barrett et al., 2020; Deehan et al., 2015; Doyon et al., 2013; Lárraga et al., 2017). Variations also exist in nicotine administration across these studies. Some studies utilized noncontingent consumption through drinking solutions (Lallemand et al., 2007; Marshall et al., 2003; Potthoff et al., 1983), others used contingent consumption (Deehan et al., 2015), or noncontingent systemic injections (Barrett et al., 2020; Doyon et al., 2013; Smith et al., 1999). Still, others relied on contingent intravenous self-administration protocols (Chandler et al., 2020; Lárraga et al., 2017). These preclinical studies present mixed results, with some suggesting that nicotine enhances the reinforcing effects of alcohol (Barrett et al., 2020; Doyon et al., 2013; Lallemand et al., 2007; Lárraga et al., 2017; Marshall et al., 2003; Smith et al., 1999), while others show no significant effect (Chandler et al., 2020; Deehan et al., 2015; Marshall et al., 2003). Notably, in the two instances where rats self-administered both nicotine and alcohol, both substances were simultaneously delivered, either in an oral solution (Deehan et al., 2015) or through an intravenous infusion (Lárraga et al., 2017). It is important to acknowledge that significant challenges exist in replicating concurrent substance use in preclinical studies due to limitations in technology, equipment, and current scientific understanding. Thus, there is a critical knowledge gap concerning the interaction of nicotine and ethanol when both substances are independently self-administered using translationally relevant routes of administration, specifically intravenous for nicotine and oral for alcohol.

Individuals diagnosed with alcohol use disorder are substantially more likely to smoke cigarettes compared to occasional drinkers. Notably, estimates suggest that 80-90 % of people with alcohol use disorder also engage in smoking. The smoking rate in this population is considerably higher than in those not diagnosed with alcohol use disorder (Burling and Ziff, 1988; DiFranza and Guerrera, 1990; Toneatto et al., 1995). Research has demonstrated that the consumption of ethanol can markedly increase cigarette smoking among individuals with alcohol use disorders. In contrast, ethanol does not affect cigarette consumption in those without an alcohol dependency (Henningfield and Goldberg, 1983; Mintz et al., 1985).

Additional reports indicate that ethanol may increase cigarette smoking in regular alcohol consumers (consuming 4-10 drinks a week) who are also moderate-to-heavy smokers (smoking 20-30 cigarettes a day; Mitchell et al., 1995). Contrary to the wealth of clinical literature detailing the effects of alcohol on cigarette smoking, there are scarcely any parallel reports in the preclinical field. A few relevant studies suggest that ethanol can diminish nicotine’s discriminative cues in a two-lever discrimination paradigm (Korkosz et al., 2005; although see Le Foll and Goldberg, 2005). Some findings also indicate that a combined nicotine and ethanol stimulus can produce a discriminative cue distinct from those evoked by either substance alone (Troisi et al., 2013). In summary, existing findings imply that ethanol may increase the reinforcing effects of nicotine. Nevertheless, a broader array of comprehensive preclinical investigations is essential for deepening our understanding of this complex interaction.

One of the programmatic ways to assess the relationship between ethanol and nicotine reinforcement is with the help of reinforcer demand modeling. Originally adapted from microeconomic theory, this method links the consumption of goods to expenditure, and it is used extensively to study behavioral responses maintained by diverse reinforcers in both clinical and preclinical settings. This method has been broadly applied to evaluate behavioral responses driven by a range of factors, such as sensory stimulation, food, and drugs, among others (Hursh et al., 2005; Hursh and Roma, 2016). In this methodology, rats can be trained to respond for a reinforcer on a low fixed ratio (FR) schedule of reinforcement. Over successive sessions, this ratio is incrementally increased, thereby raising the “cost” of obtaining a reinforcer. The reinforcer is conceptualized as a “good,” the response output maintained by the reinforcer as “consumption expenditure,” and the FR schedule as “cost.” Reinforcer demand modeling generates rich, grouped, and individual data, facilitating the examination of various aspects of behavior related to reinforcement in the particular context or a particular experimental design (Killeen and Jacobs, 2017). It can provide demand indices to describe demand at a price of zero (simulating free availability), demand elasticity, maximum expenditure, and the price at which demand becomes inelastic. Importantly, it allows for the assessment of the reinforcer’s strength, represented by the degree to which a subject is willing to work for a reinforcer, a concept known as the essential value (EV). The advantage of using the essential value is that it offers a standardized measure across different commodities and enables the comparison of these commodities across the reinforcement spectrum. For instance, it can be used to compare the reinforcing value of heroin to that of cocaine, benzodiazepines, or even chicken wings (Schwartz et al., 2021). We previously used this approach to assess grouped and individual economic demands for substances like heroin and nicotine (Kazan et al., 2020; Kazan and Charntikov, 2019; Stafford et al., 2019). The ability to evaluate individual preferences for different reinforcers and subsequently apply predictive modeling to the gathered data makes the reinforcer demand modeling a highly suitable approach for the investigation of the interactions between nicotine and ethanol.

The present study was designed to systematically evaluate the interactions between nicotine and ethanol, employing a model that permits concurrent self-administration of both substances by rats in a translationally relevant, long-access (12-hour) setting. The design aimed to mimic extensive daily substance consumption patterns where rats voluntarily intake each substance for half a day before abstaining for the remaining duration. To capture relevant data, we chose a within-subjects design that treats substance consumption as a continuous variable, thereby eliminating the need for substance-abstaining controls. This design approach enables the assessment of economic demands for various substances in the same population of subjects, facilitating the comparison of indices derived from those demand models using predictive modeling. It also enables the investigation of whether rats with a high economic demand for nicotine exhibit a correspondingly high demand for ethanol. Additionally, it examines if specific indices of ethanol demand can predict those derived from nicotine demand. Further, the indices derived from the economic demand for each factor, considered separately, can be used to predict responses when both substances are simultaneously self-administered (concurrent self-administration). Importantly, this model of concurrent nicotine and ethanol self-administration can reflect conditions when one substance is available at a relatively low price (FR1 schedule of reinforcement) or a higher price (progressive ratio schedule of reinforcement; PR). Additionally, such study design can include sessions where a secondary substance is administered noncontingently to further evaluate its impact on the primary substance. In summary, this study has been designed to examine the interplay between ethanol and nicotine, utilizing a model that closely parallels clinical usage. This design enables a comprehensive evaluation of individual data collected from the self-administration of each substance alone, as well as their concurrent use.

The main objective of this study was to investigate nicotine and ethanol interactions using a relevant self-administration model for each substance. However, these types of studies also present an opportune setting to collect and analyze additional, pertinent data. One instance of this is the evaluation of both sucrose and sweetened ethanol economic demands, as facilitated by the process of sucrose fading to establish ethanol self-administration. These supplementary data can potentially shed light on whether an individual’s preference for ethanol is linked to a preference for primary rewards such as sucrose or if the sweetened ethanol consumption is driven by the added sucrose. This data holds the potential to meaningfully enhance our understanding of the interactions between primary rewards and ethanol. Further validation for the economic demand for ethanol can be attained by assessing the withdrawal symptoms from ethanol. In theory, if a rat shows higher demand for ethanol, it might also exhibit stronger withdrawal symptoms. This could serve as a validation for using economic demand modeling as a tool for identifying susceptibility to ethanol use (represented by higher economic demand). The inclusion of such data can further enhance our holistic understanding of the study’s results. In summary, this study provides a novel methodical approach to exploring the interaction between ethanol and nicotine use, while collecting a wider array of individual data to facilitate a deeper understanding of their interactions.

## 2. Materials

### 2.1. Subjects

Twenty-two male Wistar rats weighing between 250-300 g were obtained from Envigo (Indianapolis, IN, USA). The rats were individually housed in a temperature-controlled vivarium with a 12-hour light/dark cycle, with lights turning on at 0700. After being introduced to the colony, the rats were allowed to acclimate for one week before the commencement of the experimental procedures. During this acclimation period and for one week after the implantation of the intravenous catheter, the rats had access to food and water ad libitum. Throughout the study, the rats were subjected to food restriction in order to maintain their weight at 90 % of their free-feeding weight, with water available without restriction. The free-feeding weight was gradually increased by 2 g every 30 days. All procedures were conducted in accordance with the Guide for the Care and Use of Laboratory Animals (National Research Council et al., 2010) and were reviewed and approved by the University of New Hampshire Institutional Animal Care and Use Committee.

### 2.2. Apparatus

#### 2.1.1. Self-administration chambers

Behavioral tests were conducted in sound- and light-attenuated Med Associates conditioning chambers (30.5 × 24.1 × 21.0 cm; l × w × h) equipped with an exhaust fan (ENV-018MD; Med Associates, Inc.; St. Albans, VT, USA). The chambers had aluminum sidewalls, metal rod floors, and polycarbonate surfaces. Two retractable levers (147 nN required for micro-switch closure) were mounted on each side of the right-side wall and were used as manipulanda to operate the retractable sipper equipped with a lickometer (ENV-252M; Med Associates, Inc.; St. Albans, VT, USA) positioned on the wall between those levers. Cue lights were positioned above each lever. For nicotine self-administration, two nosepokes with a yellow LED and an infrared beam monitoring the entry were installed on the sidewall opposite the levers. The infusion pump (PMH-100VS; Med Associates; St. Albans, VT, USA) for each chamber was located outside the sound-attenuating cubicle. A 5 mL syringe mounted on the infusion pump was connected to a swivel coupled with a spring leash (C313C; Plastics One; Roanoke, VA, USA) and Tygon® tubing (AAQ04103; VWR; West Chester, PA, USA) suspended over the chamber’s ceiling on a balanced metal arm. For nicotine-alone self-administration, levers and retractable sippers were removed from the chamber. Med Associates interface and software (Med-PC for Windows, version IV) were used to collect data and execute programmed events.

#### 2.2.2. Open Field

Open-field tests were conducted in an open-top square plywood box (120 cm × 120 cm × 25 cm; l × w × h) painted with flat black enamel. Test sessions were recorded by a camera mounted above the apparatus and processed using the ANY-maze video tracking system (Stoelting Co.; Wood Dale, IL, USA).

#### 2.2.3. Elevated Plus-Maze

Elevated plus-maze tests were conducted using the elevated plus-shaped platform (Stoelting Co.; Wood Dale, IL, USA; lane width = 10 cm, arm length = 50 cm, wall height = 40 cm, leg height = 40 cm). Test sessions were recorded by a camera mounted above the apparatus and processed using the ANY-maze video tracking system (Stoelting Co.; Wood Dale, IL, USA).

### 2.3. Drugs

Ethanol (200 proof; Decon Labs; King of Prussia, PA, USA) and sucrose (store-bought sugar) solutions were made using tap water. Nicotine bitartrate (MP Biomedicals; Solon, OH, USA) was dissolved in 0.9 % sterile saline. The pH of nicotine was adjusted to 7.0 ± 0.2 with a dilute NaOH solution. Nicotine doses are reported as a base. Doses and administration protocols were adopted from previous research (Charntikov et al., 2021; Kazan et al., 2020).

## 3. Methods

Figure 1 presents the experimental progression. Before testing, all rats underwent a three-day period of twice-daily handling by all experimenters. Baseline behavioral assessments were performed using elevated plus-maze and open field tests. Following the baseline assessments, rats were trained to lever press for a liquid reward and then evaluated for sucrose, sweetened ethanol, ethanol-alone, and nicotine economic demands in sequential order. Blood ethanol concentration tests and reassessment of behaviors using elevated plus-maze and open field tests occurred after the ethanol-alone demand assessment and before the nicotine demand assessment. Rats were then subjected to a co-administration experiment to determine the effect of one substance on the self-administration of another substance under different schedules of reinforcement and contingencies. Detailed experimental methods are described below.

**Fig. 1.**
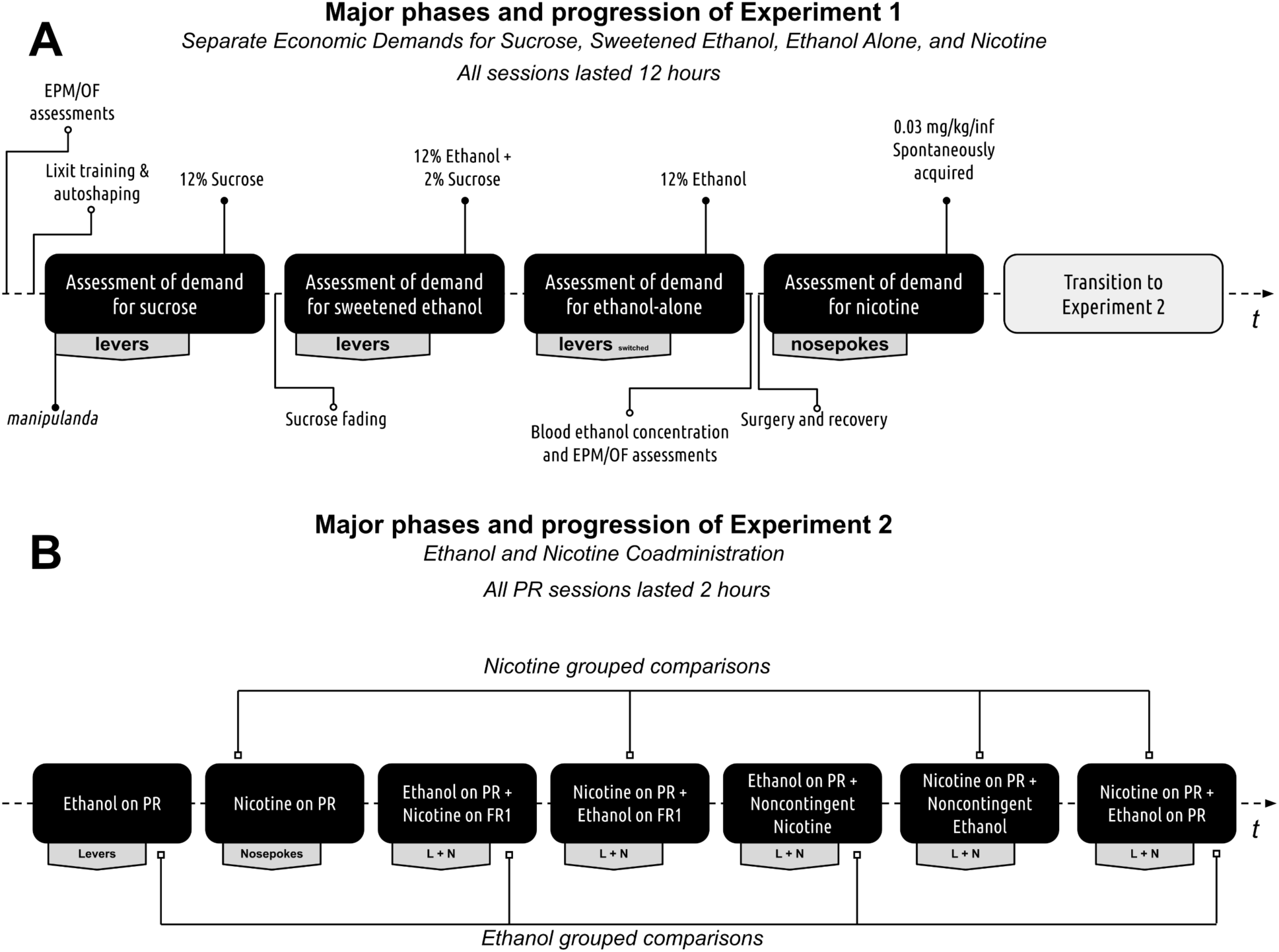
Experimental Progression. (A) Progression of Experiment 1. Rats initially underwent assessments using the elevated plus maze and open field tests. Following this, they were trained to retrieve rewards from a retractable sipper tube and then trained to self-administer a 12 % sucrose solution. The rats’ individual demands for sucrose, sweetened ethanol, ethanol-alone, and nicotine were then evaluated. Before the nicotine self-administration phase, the rats were reassessed on the elevated plus maze and open field tests during withdrawal. Finally, blood ethanol concentration was measured at the conclusion of the ethanol self-administration phase to confirm ethanol consumption. (B) Progression of Experiment 2. All rats were initially evaluated for baseline responses on the Progressive Ratio (PR) schedule of reinforcement. Following this, their performance was assessed during ethanol and nicotine co-administration, with one substance restricted to a PR schedule of reinforcement while the other was available on a Fixed Ratio 1 (FR1) schedule, a PR schedule, or administered non-contingently.

### 3.1. Experiment 1

#### 3.1.1. Open field test

Rats were first acclimated to the testing room for 60 min in their home cages. Subsequently, they were placed individually into the center of the open field apparatus for 10 min and then returned to the vivarium. The open-field apparatus was divided into two portions: the center consisted of a central 60 cm x 60 cm square (located 30 cm from the apparatus wall), while the remaining surrounding area of the apparatus consisted of the perimeter. The ANY-maze software was used to collect the total time spent in the center of the open field, total distance traveled, average travel speed, the total number of freezing episodes, and total freezing time. Dependent measures were divided into the first 5 min (habituation; 0-5 min) and the last 5 min (test; 5-10 min) of the test, with behaviors during the second 5 min bin used for data analyses. Open field tests were conducted before the lever training (see Figure 1) and during ethanol withdrawal. Withdrawal tests were performed after the acquisition of ethanol-alone economic demand (10-11 hours after the end of the ethanol-alone self-administration session).

#### 3.1.2. Elevated plus-maze test

Before testing, rats were acclimated for 60 minutes in their home cages. Subsequently, rats were individually placed in the center of the elevated plus-maze apparatus for a 10-minute test session, after which they were returned to the vivarium. Data, including total distance traveled, average travel speed, total number of freezing episodes, total freezing time, and total time in the open arms, were collected using ANY-maze software. Dependent measures were divided into the first 5 minutes (habituation; 0-5 min) and the last 5 minutes (test; 5-10 min) of the test. The second 5-minute bin was used for data analysis. The elevated plus-maze tests were performed before the lever training (as depicted in Figure 1) and during withdrawal from ethanol. Withdrawal tests were conducted after the acquisition of ethanol-alone economic demand, 10-11 hours after the end of the ethanol-alone self-administration session.

#### 3.1.3. Preliminary lever training

Rats were first trained to consume sucrose (12 % w/v) from a retractable sipper. These sipper training sessions lasted 120 min, during which non-contingent sucrose presentations were delivered on a variable time interval (∼ 3 rewards per minute). Rats were then trained to lever-press for the 12 % sucrose solution using an auto-shaping procedure. Each session began with the illumination of the house light and the insertion of a randomly selected lever (right or left). Lever presses or a lapse of 15 s resulted in the insertion of a sipper tube, lever retraction, extinction of the house light, and the illumination of cue lights located above each lever. Fifteen seconds later, the sipper tube was retracted, cue lights turned off, and the house light was turned on, following which a randomly selected lever was inserted back into the chamber. The same lever could not be presented more than twice in a row, and the number of left and right lever presentations was equal across the session. Training continued until rats made lever presses on at least 80 % of lever insertions for two consecutive days (total training time was 3-6 daily sessions based on individual performance). One rat was excluded from the study due to an inability to acquire lever-pressing behavior.

#### 3.1.4. Acquisition of economic demand for 12 % sucrose

Rats were assigned active levers pseudo-randomly with the condition that there was an equal number of right and left active levers. Rats were then trained to self-administer 12 % sucrose on a fixed schedule of reinforcement (FR1) for three consecutive days. Each session began with the insertion of both levers and the illumination of a house light. When the schedule requirement was reached, a sipper tube was inserted into the chamber, the levers were retracted, and cue lights were illuminated. Five seconds later, the sipper tube was retracted, the levers were reinserted, the cue lights were turned off, the house light was turned on, and the rats were able to continue pressing the levers for a liquid reward. Self-administration sessions were conducted during the night cycle, which corresponds to the rodents’ active phase (1900-0700). After three days of 12 % sucrose self-administration, the rats earned sucrose on a fixed ratio (FR) schedule of reinforcement that was escalated daily (between-sessions escalation) using the following sequence: 1, 3, 5, 8, 12, 18, 26, 38, 58, 86, 130, 195, 292, 438, and 657. Rats progressed through these daily schedule escalations until failing to earn at least one reinforcer. Subsequently, rats were allowed to self-administer 12 % sucrose on a variable schedule of reinforcement (VR3; range 1-5) until all rats completed demand assessment, plus an additional three daily sessions to reacquire 12 % sucrose self-administration.

#### 3.1.5. Sucrose fading

The rats were trained to self-administer ethanol solution using a sucrose-fading procedure during 12-hour sessions. The active lever assignment remained the same as the previous phase. The procedure followed the same heuristics as the previous phase, but the liquid reinforcer was adjusted. At the beginning of the training, the rats were given 12 % sucrose solution, to which progressively higher ethanol concentrations were added every four days using the following sequence: 2 %, 4 %, 8 %, and 12 %. The rats were allowed to self-administer 12 % sucrose and 12 % ethanol solution for 6 consecutive days, and then the sucrose concentration was gradually decreased to 2 % using the following sequence: 12 %, 8 %, 4 %, and 2 % (four days per each concentration). After the fading protocol, rats self-administered 2 % sucrose and 12 % ethanol solution using the VR3 schedule of reinforcement.

#### 3.1.6. Acquisition of economic demand for sweetened ethanol (2 % sucrose and 12 % ethanol solution)

The acquisition of economic demand for sweetened ethanol followed the same procedure as that for 12 % sucrose, but with 2 % sucrose and 12 % ethanol solution as the reinforcer. Afterward, rats underwent self-administration of 2 % sucrose and 12 % ethanol solution on a VR3 schedule of reinforcement until all rats completed demand assessment and an additional three daily sessions to reacquire self-administration behavior.

#### 3.1.7. Acquisition of economic demand for ethanol-alone

The 2 % sucrose and 12 % ethanol solution was replaced with a 12 % ethanol-alone solution. Active and inactive lever assignments were reversed, and the cues associated with access to ethanol were changed to avoid any possible confounding effects of conditioned reinforcement associated with the sucrose reward. Each ethanol-alone self-administration session began with both levers inserted and both cue lights illuminated. Upon reaching the schedule requirement a sipper tube was inserted, levers were retracted, cue lights were turned off, and the house light was illuminated. The sipper tube was retracted five seconds later, cue lights were turned on, the house light was turned off, and rats could continue lever pressing for ethanol. Using this protocol, rats self-administered ethanol-alone on a VR3 schedule of reinforcement for 7 to 10 daily 12-hour sessions until the number of active lever presses exceeded the number of inactive lever presses. The acquisition of economic demand for ethanol-alone followed and was identical to the protocol described earlier with 12 % sucrose. Once the terminal schedule requirement was reached, where rats failed to earn at least one reinforcer, all rats were allowed to self-administer ethanol-alone on a VR3 schedule of reinforcement until all rats completed demand assessment plus an additional three daily sessions to reacquire ethanol self-administration. All ethanol self-administration sessions were conducted during the night cycle and lasted for 12 hours.

#### 3.1.8. Plasma ethanol concentration tests

After the acquisition of economic demand for ethanol-alone, rats self-administered ethanol on a VR3 schedule of reinforcement until the completion of open field, elevated plus-maze, and plasma ethanol concentration tests, which were separated by at least two daily ethanol-alone self-administration sessions. Blood ethanol concentration tests occurred immediately after one hour of ethanol-alone self-administration that substituted a regular 12-hour session. There were two plasma alcohol concentration tests separated by at least two days of ethanol-alone self-administration. To collect plasma samples, rats were lightly restrained in a towel, and approximately 300 microliters of blood was collected via lateral tail vein incision while the tail was placed into 46±2°C water to promote vasodilation. The first incision was made in the distal 2 cm of the tail, with subsequent incisions made at least 1 cm rostral to the previous. All samples were collected within 3 min, and rats were returned to a home cage within 5 min (Drugan et al., 2005; Stafford et al., 2019). The plasma samples were centrifuged at 4°C for 4 min at 1300 rpm to separate red blood cells, and the plasma was stored at -80°C until assay. An Ethanol Assay Kit (ab65343; Abcam; Cambridge, UK; McCarter et al., 2017) was used to measure the average plasma alcohol concentration from both samples for each rat.

#### 3.1.9. Catheter implantation surgery

Anesthesia was induced with 5 % isoflurane for 5 minutes and maintained at ∼2.5 % for the remainder of the surgery. Butorphanol (5 mg/kg; SC) and meloxicam (0.15 mg/kg; SC) were administered for pain management. The catheter was implanted in the right external jugular vein and routed around the ipsilateral shoulder to a polycarbonate access port (313-000B; Plastics One Inc.; Roanoke, VA, USA) implanted along the dorsal midline. Cefazolin (50 mg/mL) diluted in sterile saline with heparin (30 U/mL) was used to flush the catheter and maintain patency throughout the self-administration phase. Rats were monitored and given at least one week to recover before progressing to nicotine self-administration. After completing the nicotine self-administration phase or when suspecting catheter patency loss, catheter patency was assessed by infusing 0.05 mL xylazine (20 mg/mL) through the IV catheter. Eight rats were excluded from the study due to the loss of catheter patency throughout the study. All data from these eight rats prior to suspicion of catheter patency loss was included in the final dataset. Additional three rats did not recover from the surgery.

#### 3.1.10. Nicotine self-administration and nicotine demand

For nicotine self-administration, the chambers were reconfigured by changing the manipulanda from levers to nosepokes. This was done to reduce the conditioned enhancement of reinforcing effects associated with the previous manipulanda, which were paired with the sucrose and ethanol stimuli. Rats spontaneously acquired nicotine self-administration using nosepokes as manipulanda. The start of each session was signaled by turning on the nosepoke lights and priming the catheter with nicotine (31 µL or 90 % of internal catheter volume). The active nosepoke was initially reinforced using a VR1.5 schedule of reinforcement (range 1-3; 3-5 days) and then using a VR3 schedule of reinforcement (range 1-5; 3-5 days), with the inactive nosepoke being available but having no programmed consequence. Upon meeting the schedule requirement, rats received a ∼1-sec infusion of nicotine (0.03 mg/kg/infusion), and the nosepoke lights were extinguished for a 3-sec timeout, during which rats were unable to earn an infusion. The exact dose of nicotine was self-administered by all rats using a slight variation in infusion duration, which was automatically calculated by the program based on their pre-session weight. All nicotine self-administration sessions lasted for 12 hours and were conducted during the night cycle. The acquisition of economic demand for nicotine was identical to the acquisition of economic demand for ethanol, except that the reinforcer was nicotine. After reaching a terminal schedule requirement, where rats failed to earn at least one reinforcer per session, all rats were allowed to self-administer nicotine-alone on a VR3 schedule of reinforcement until all rats completed demand assessment plus an additional 3 daily sessions to reacquire nicotine self-administration.

### 3.2. Experiment 2

#### 3.2.1. Concurrent ethanol and nicotine self-administration

Given the impracticality of maintaining catheter patency throughout the duration of concurrent ethanol and nicotine self-administration tests using a behavioral economics approach that involves between-session price escalation, we opted to use the response on the progressive schedule of reinforcement as an alternative measure. This measure reflects the extent of effort rats are willing to exert for each substance individually or in the presence of a secondary substance. For establishing a performance baseline, rats were first evaluated on a progressive ratio (PR) schedule of reinforcement, where the response to each substance alone served as a baseline in comparative statistical tests. This PR schedule, identical to the between-session progression utilized for economic demand, comprised of the following sequence: 1, 3, 5, 8, 12, 18, 26, 38, 58, 86, 130, 195, 292, 438, and 657. During these sessions, each rat self-administered a substance for two hours on a PR schedule, followed by another ten hours on a VR3 schedule for the same reinforcer. This two-hour limit aimed to constrain learning about nonreinforcement. Subsequently, rats were allowed to self-administer both substances simultaneously. Both levers and nosepoke manipulanda were available for concurrent self-administration of ethanol and nicotine. Each concurrent self-administration session lasted 12 hours. In the initial two-hour segment of the session, rats earned a primary substance using the PR reinforcement schedule, while a secondary substance was concurrently available either on an FR1 or a PR reinforcement schedule. To limit learning about non-reinforcement, in the remaining ten-hour segment of the session, rats self-administered the primary substance using a VR3 schedule, while the secondary substance was available on the FR1 schedule. Separate two-hour control sessions were conducted wherein the primary substance was self-administered on a PR reinforcement schedule, while the secondary substance was noncontingently administered (nicotine) or presented (ethanol). During these control sessions, noncontingent nicotine infusions or access to ethanol occurred within pre-established parameters and alongside previously identified associated cues. These single noncontingent infusions or ethanol deliveries took place at the start of the session and every 10 minutes thereafter.

Figure 1B shows the progression of this experimental phase and outlines the structure of ethanol and nicotine coadministration sessions. There was a total of five sessions for each testing combination. The first three sessions were conceptualized as an acclimation to the protocol and schedule conditions. Data from the last two testing sessions for each combination were used for statistical analyses. In this experimental phase, concurrent self-administration sessions for ethanol and nicotine were conducted using different combinations of primary and secondary substances with different schedules of reinforcement. The testing combinations and accompanying schedules of reinforcement are presented in Table 1. This experimental design allowed the sampling of baseline responding on the progressive ratio (PR) schedule for each substance alone, sampling of responding on the PR schedule for a primary substance while a secondary substance was available either at a low cost (FR1) or at a relatively high cost (PR), and sampling of responding on the PR schedule of reinforcement while the secondary substance was administered non-contingently.

**Table 1:** Ethanol and Nicotine Coadministration Testing Combinations and Schedules of Reinforcement.

| **Primary Substance** | **Reinforcement Schedule** | **Secondary Substance** | **Reinforcement Schedule** |
| --- | --- | --- | --- |
| Nicotine | PR | N/A | N/A |
| Nicotine | PR | Ethanol | FR1 |
| Nicotine | PR | Ethanol | Noncontingent |
| Nicotine | PR | Ethanol | PR |
| Ethanol | PR | N/A | N/A |
| Ethanol | PR | Nicotine | FR1 |
| Ethanol | PR | Nicotine | Noncontingent |

PR - progressive ratio schedule of reinforcement, FR1 - fixed ratio 1 schedule of reinforcement, N/A - not available

Our study’s co-administration procedures address several key aspects of ethanol and nicotine use. First, concurrent self-administration of both substances attempts to simulate various co-use scenarios, capturing the often-overlooked dynamics and interactions between ethanol and nicotine use. Second, varied reinforcement schedules help investigate the impact of effort on substance use, revealing how accessibility and cost influence behavior, as seen through fixed (FR) and progressive (PR) ratio schedules. Third, progressive ratio schedules enable us to measure the effort rats exert to self-administer substances, avoiding lengthy economic demand assessments that could compromise intravenous catheter functionality. Lastly, non-contingent administration of secondary substances allows us to study their influence on reinforcing effects of primary substances, providing a holistic perspective on multi-substance use behavior. While our current methodology doesn’t extend to exploring areas such as cross-commodity demand and cross-price elasticity—potential avenues for future research—it does afford a nuanced and comprehensive approach to studying the complexities of substance use behavior. This methodology integrates various facets of the addiction cycle, which are often explored in isolation, facilitating a more cohesive understanding.

### 3.3. Data analysis

#### 3.3.1. Economic demand

The economic demand for a reinforcer was assessed using Hursh and Silberberg’s operant demand framework (Hursh, 2014; Hursh and Silberberg, 2008). Consumption data (g/kg) from each reinforcement schedule were fit into the nonlinear least squares regression model using the formula: 𝑙𝑜𝑔𝑄 = 𝑙𝑜𝑔𝑄0 + 𝑘 × (*e*^(−α*Q*0*C*)^ − 1). Here, *Q* denotes quantity consumed, *Q_0_* indicates quantity consumed when the price is zero (i.e., consumption at zero cost or maximal consumption), 𝑘 is a parameter that adjusts the range of the dependent variable (*logQ*), *e* is the base of the natural logarithm, *C* is the cost, and α is the rate of decline in consumption as cost increases (demand elasticity). The model estimated the demand elasticity (*α*) and intensity (*Q_0_*). Maximum expenditure (*O_max_*) was calculated using the highest expenditure for each price or reinforcement schedule. The point of price where demand becomes elastic and expenditure reaches maximum (*O_max_*) is represented by *P_max_*. The Essential Value (*EV*), calculated as 1/(100 × α × k ^1.5^), inversely proportional to *α*, was derived from the economic demand model. *EV* quantifies a reinforcer’s ability to maintain operant behavior amidst escalating behavioral costs and is often used to signify the intensity of demand or the value of a commodity. The consumption values were initially log-transformed, and then the economic demand was derived from those values using a least-squares nonlinear fit via GraphPad Prism version 9 (GraphPad Software, Inc., La Jolla, CA).

All other statistical analyses were conducted in R 4.1.3 (R Core Team, 2019). Variables, including blood ethanol concentration, economic demand, and concurrent self-administration comparisons, were assessed using linear mixed-effects modeling via the {nlme} package for R (Pinheiro et al., 2017). This analysis method, preferred over ANOVA, doesn’t necessitate homogeneity or independence of data cases, can model interrelated outcomes, and is robust in handling missing data or unequal group sizes—common in preclinical animal models. Demand indices across various substances and stress indices’ association with ethanol demand were examined using simple regression analyses. Effects of ethanol withdrawal on behavioral outcomes were assessed via paired samples t-tests using data from the elevated plus maze and open field tests. In our supplemental analyses, we used simple linear regressions to explore how well economic demand parameters predict PR schedule reinforcement response, with total active lever presses and demand parameters as dependent and independent variables, respectively. Furthermore, we conducted multiple linear regressions, including all demand parameters, for a general comparison in all our supplemental analyses.

In our concurrent ethanol and nicotine self-administration tests, we used total active lever presses as a primary dependent measure. Instead of relying on the commonly used breaking point derived from the progressive ratio (PR) schedule of reinforcement, we chose to utilize active lever responding due to its numerous advantages. Firstly, active lever responding enhances sensitivity and enables the detection of subtle behavioral differences between treatment conditions, even when variations are small. Secondly, considering the entire response profile of active lever responding improves statistical power by increasing the number of observations for statistical analysis, thereby enhancing the reliability of our findings. Lastly, by aligning our research more closely with the assessment of operant behavior in humans, active lever responding increases the translational relevance of our findings. Consequently, active lever responding was selected as the primary dependent measure to investigate the interaction effects in our co-administration tests.

## 4. Results

### 4.1. Blood ethanol concentration

The volume of consumed ethanol predicted blood ethanol concentration (*χ*^2^_(1)_ = 10.48, *p* = 0.0012) and explained 22 % in blood ethanol concentration variance (*R*^2^ = 0.22). This confirms a positive relationship between ethanol volume consumed and blood concentration, validating its reliability as a measure of consumption in this context.

### 4.2. Comparing demand for sucrose, sweetened ethanol, and ethanol-alone

Figure 2 illustrates the demand curves for each substance being investigated (first column). For comparison, this figure also presents the individual highest and lowest demand curves for each substance (second column). In addition, Figure 2 shows the essential values for each substance, highlighting the variability of essential values across substances (third column). 𝐸𝑉 differed significantly between self-administered substances (χ^2^_(3)_ = 70.29, 𝑝 < .0001/. The essential value for sucrose was significantly higher than the essential values for sweetened ethanol, ethanol-alone, or nicotine (Figure 3A). The elasticity of demand, represented by 𝛼 value, also differed significantly across self-administered substances (χ^2^_(3)_ = 13.04, 𝑝 < .01/. The elasticity of demand for ethanol-alone surpassed that of sucrose, sweetened ethanol, or nicotine (Figure 3B), indicating a higher sensitivity to changes in cost for ethanol. Significant differences were observed in the initial level of consumption (𝑄_%_) across self-administered substances (χ^2^_(3)_ = 93.67, 𝑝 < .0001/, with sucrose eliciting a higher initial level of demand compared to sweetened ethanol, ethanol-alone, or nicotine (Figure 3C). Moreover, the maximum output consumption level (𝑂_,-._) varied significantly between self-administered substances (χ^2^_(3)_ = 71.46, 𝑝 < .0001/. 𝑂_,-._ for sucrose was significantly higher than 𝑂_,-._ for sweetened ethanol, ethanol-alone, or nicotine (Figure 3D). The maximum price paid (𝑃_,-._) also differed significantly between self-administered substances (χ^2^_(3)_ = 13.97, 𝑝 < .01/. 𝑃_,-._ for sucrose was significantly higher than 𝑃_,-._ for ethanol (Figure 3E). Overall, these findings suggest that rats exerted greater effort (higher *EV*) for sucrose compared to the other substances. Sucrose also evoked the highest base level of demand (𝑄_%_) and maximum consumption level (𝑂_,-._). Conversely, rats demonstrated the greatest sensitivity to changes in the cost of ethanol (highest *α*). Peak response (𝑃_,-._) was highest for sucrose and lowest for ethanol, indicating a differential response pattern based on substance type.

**Fig. 2.**
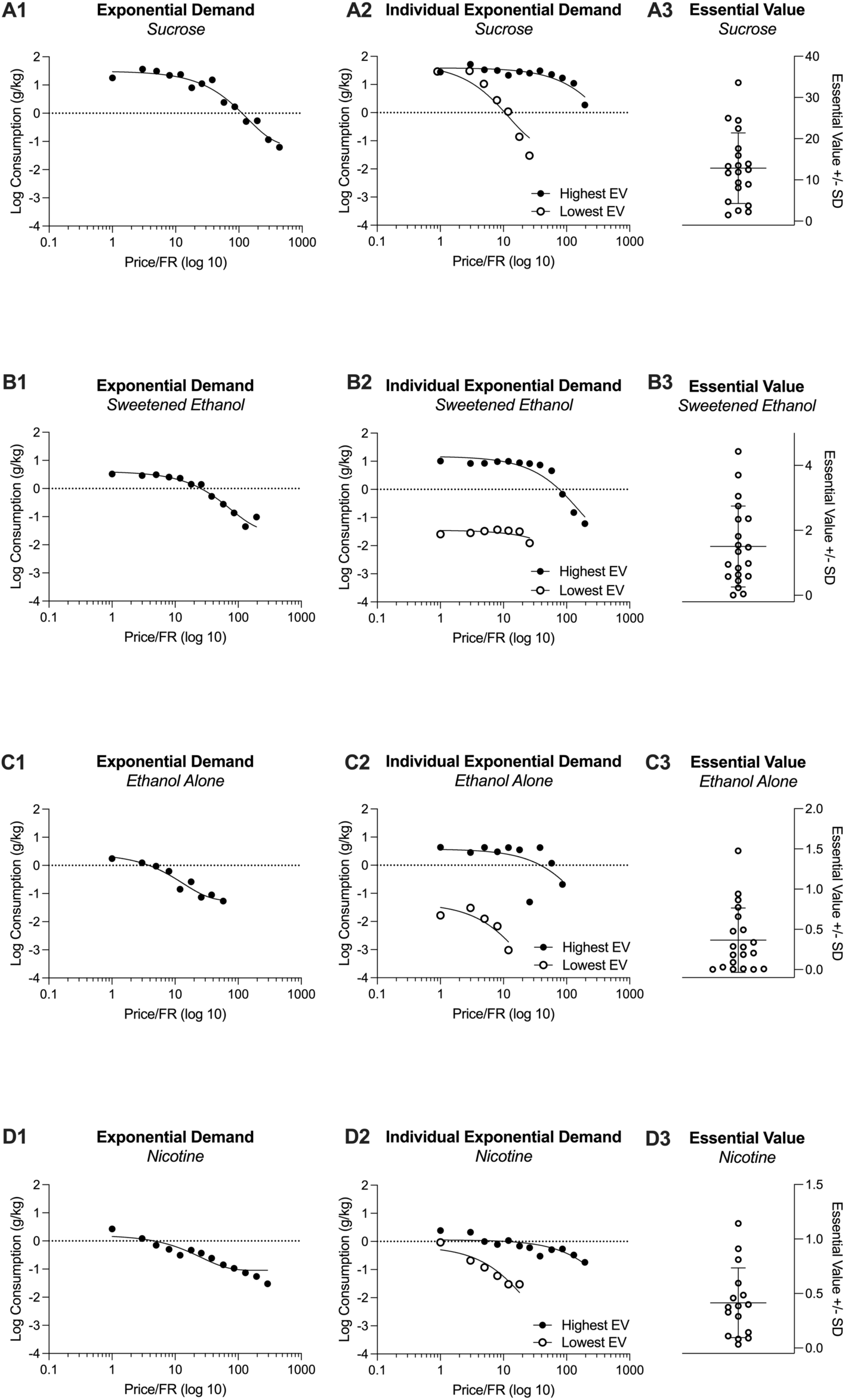
Grouped demand curves for sucrose, sweetened ethanol, ethanol-alone, and nicotine (A1, B1, C1, and D1). Individual high and low economic demand curves for each substance (A2, B2, C2, and D2). Individual Essential Values for each substance (A3, B3, C3, D3).

**Fig. 3.**
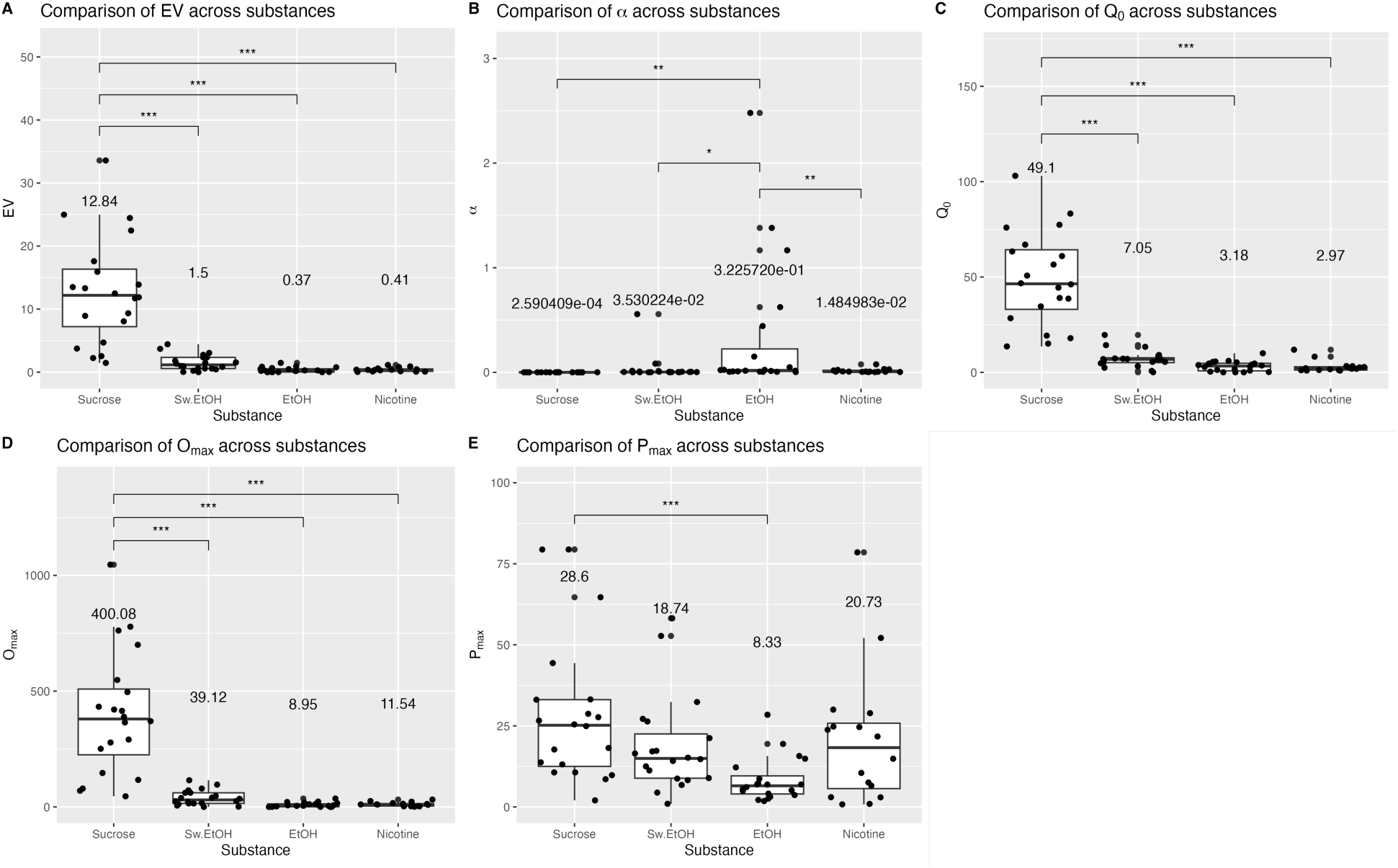
Comparison of main indices derived from the reinforcer demand modeling. Each panel’s mean value for each substance is shown above the box. In panel B, the numbers are represented in scientific notation (e.g., 3.530224e-02), which signifies a small decimal number. Here, “e-02” means that the decimal point in 3.530224 is moved two places to the left, giving 0.03530224. Similar interpretations apply for “e-01” through “e-04”.

### 4.3. Relationship of demand indices across economic demands

Table 2 outlines the statistical outcomes from all the analyses performed in this section. The Essential Value (*EV*), the elasticity of demand (*α*), and the maximum price paid 𝑃_,-._ of sucrose were found to significantly predict the corresponding parameters for sweetened ethanol. This implies that the effort rats exerted for sucrose was paralleled by their efforts for sweetened ethanol. In addition, a shared sensitivity was observed in the rats’ responses to price increases for sweetened ethanol and ethanol-alone, as represented by the *α* value. This suggests that the elasticity of demand for these two substances was interconnected. Notably, a similar sensitivity to cost changes was also found between ethanol and nicotine, indicating a potential relationship in price response patterns across these substances. It’s important to note that while these results elucidate some relationships between substance demand indices, there remains a considerable amount of unexplained variance, as indicated by the R^2^ values. This suggests that additional factors not included in the current analysis may influence substance demand, emphasizing the complex nature of substance use behavior.

**Table 2.** Prediction of demand indices across economic demands.

| Demand Indices |  | Independent Model |  |  |  |
| --- | --- | --- | --- | --- | --- |
| Predictor | DV | $\beta$ | R<sup>2</sup> | F(df) | p-value |
| **EV** |  |  |  |  |  |
| Sucrose | Sw.EtOH | 0.66 | 0.44 | F(1,17)=13.43 | **<0.01** |
| Sucrose | EtOH | -0.01 | <0.01 | F(1,17)=0.0034 | 0.95 |
| Sucrose | Nicotine | -0.42 | 0.18 | F(1,13)=2.83 | 0.12 |
| Sw.EtOH | EtOH | 0.02 | <0.01 | F(1,17)=0.0047 | 0.95 |
| Sw.EtOH | Nicotine | 0.03 | <0.01 | F(1,13)=0.0084 | 0.93 |
| EtOH | Nicotine | -0.28 | 0.08 | F(1,13)=1.10 | 0.31 |
| **$\alpha$** |  |  |  |  |  |
| Sucrose | Sw.EtOH | 0.07 | 0.018 | F(1,17)=4.484 | **0.048** |
| Sucrose | EtOH | -0.02 | 0.025 | F(1,17)=0.7155 | 0.41 |
| Sucrose | Nicotine | -0.02 | 0.075 | F(1,13)=0.1077 | 0.75 |
| Sw.EtOH | EtOH | 0.04 | <0.01 | F(1,17)=21.13 | **<0.01** |
| Sw.EtOH | Nicotine | 0.00 | <0.01 | F(1,13)=0.1425 | 0.71 |
| EtOH | Nicotine | -0.03 | <0.01 | F(1,13)=19.58 | **<0.01** |
| **Q<sub>0</sub>** |  |  |  |  |  |
| Sucrose | Sw.EtOH | 0.40 | 0.15 | F(1,17)=3.22 | 0.09 |
| Sucrose | EtOH | -0.05 | <0.01 | F(1,17)=4.44 | 0.05 |
| Sucrose | Nicotine | -0.02 | <0.01 | F(1,13)=0.36 | 0.56 |
| Sw.EtOH | EtOH | 0.11 | 0.04 | F(1,17)=0.65 | 0.43 |
| Sw.EtOH | Nicotine | -0.41 | 0.14 | F(1,13)=2.67 | 0.13 |
| EtOH | Nicotine | 0.09 | <0.01 | F(1,13)=0.10 | 0.76 |
| **O<sub>max</sub>** |  |  |  |  |  |
| Sucrose | Sw.EtOH | 0.39 | 0.15 | F(1,17)=3.07 | 0.10 |
| Sucrose | EtOH | 0.20 | 0.04 | F(1,17)=0.68 | 0.42 |
| Sucrose | Nicotine | -0.51 | 0.26 | F(1,13)=4.52 | 0.05 |
| Sw.EtOH | EtOH | 0.30 | 0.09 | F(1,17)=1.71 | 0.21 |
| Sw.EtOH | Nicotine | -0.11 | 0.01 | F(1,13)=0.15 | 0.70 |
| EtOH | Nicotine | 0.09 | <0.01 | F(1,13)=0.10 | 0.76 |
| **P<sub>max</sub>** |  |  |  |  |  |
| Sucrose | Sw.EtOH | 0.66 | 0.44 | F(1,17)=13.43 | **<0.01** |
| Sucrose | EtOH | -0.01 | <0.01 | F(1,17)=0.0034 | 0.95 |
| Sucrose | Nicotine | -0.42 | 0.18 | F(1,13)=2.83 | 0.12 |
| Sw.EtOH | EtOH | 0.02 | <0.01 | F(1,17)=0.0047 | 0.95 |
| Sw.EtOH | Nicotine | 0.03 | <0.01 | F(1,13)=0.0084 | 0.93 |
| EtOH | Nicotine | -0.28 | 0.08 | F(1,13)=1.10 | 0.31 |

p-values in bold indicate significant effects (p<0.05).

### 4.4. The effect of withdrawal from ethanol on behavioral outcomes from the elevated plus maze and open field tests

Figure 4 compares elevated plus maze and open field performance at baseline and in withdrawal from ethanol.

**Fig. 4.**
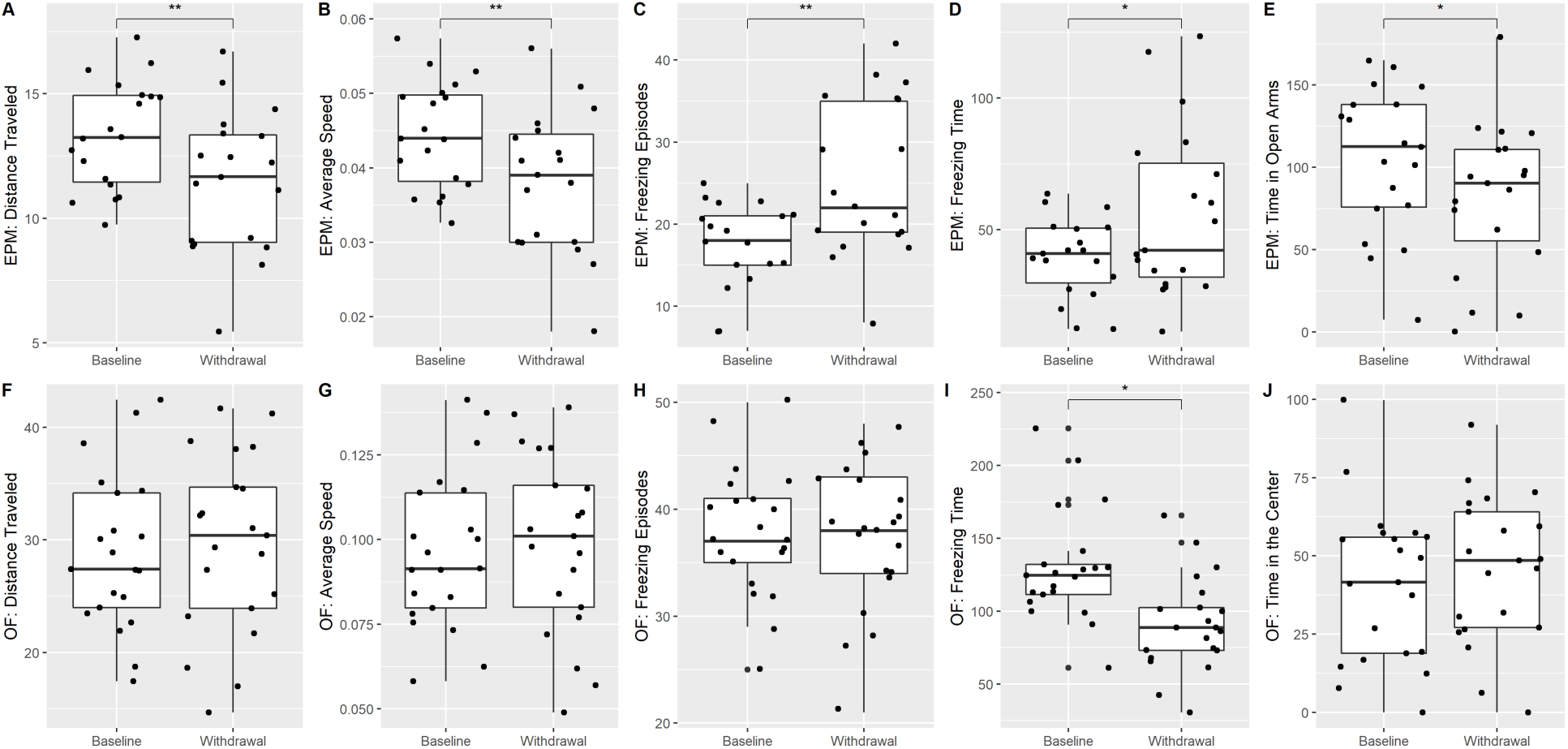
The effect of withdrawal from ethanol on behavioral outcomes from the elevated plus maze (panels A-E) and open field (panels F-J) tests. Baseline tests occurred prior to administration of any substance while withdrawal tests occurred after the acquisition of ethanol-alone economic demand.

#### Elevated Plus Maze

Distance traveled, average speed, and time spent in open arms were significantly lower during withdrawal than at baseline (𝑡(18) = 3.03, 𝑝 < 0.01; 𝑡(18) = 2.98, 𝑝 < 0.01; 𝑡(18) = 2.11, 𝑝 = 0.049; respectively). Number of freezing episodes and freezing time were significantly higher in withdrawal than at baseline (𝑡(18) = −3.25, 𝑝 < 0.01; 𝑡(18) = −2.24, 𝑝 = 0.038; respectively).

#### Open Field

Freezing time was lower during withdrawal than at baseline (𝑡(20) = 3.34, 𝑝 < 0.01).

These findings show that elevated plus maze paradigm seems to be more sensitive in detecting behavioral withdrawal effects from ethanol.

### 4.5. Association between stress indices and the demand for ethanol-alone

We utilized individual indices from the elevated plus maze and open field tests to conduct simple linear regressions. The aim was to determine whether the individual demand for ethanol, being the previously self-administered substance, could predict withdrawal symptoms as assessed by these paradigms (Table 3; Independent Model). The regression analyses revealed that the Essential Value (*EV*) of ethanol significantly predicted outcomes on four out of five indices in the elevated plus maze paradigm. These indices included distance traveled, average speed, the number of freezing episodes, and the duration of freezing time. Similarly, the *EV* of ethanol significantly predicted two out of five indices in the open field paradigm, namely distance traveled and average speed. Collectively, these results suggest that the economic demand for ethanol, as represented by its *EV*, can serve as a reliable predictor of stress-related behavioral responses during withdrawal from ethanol. The predictive capacity of *EV* in relation to these behavioral measures underscores its potential utility in understanding the complexities of ethanol withdrawal.

**Table 3.**
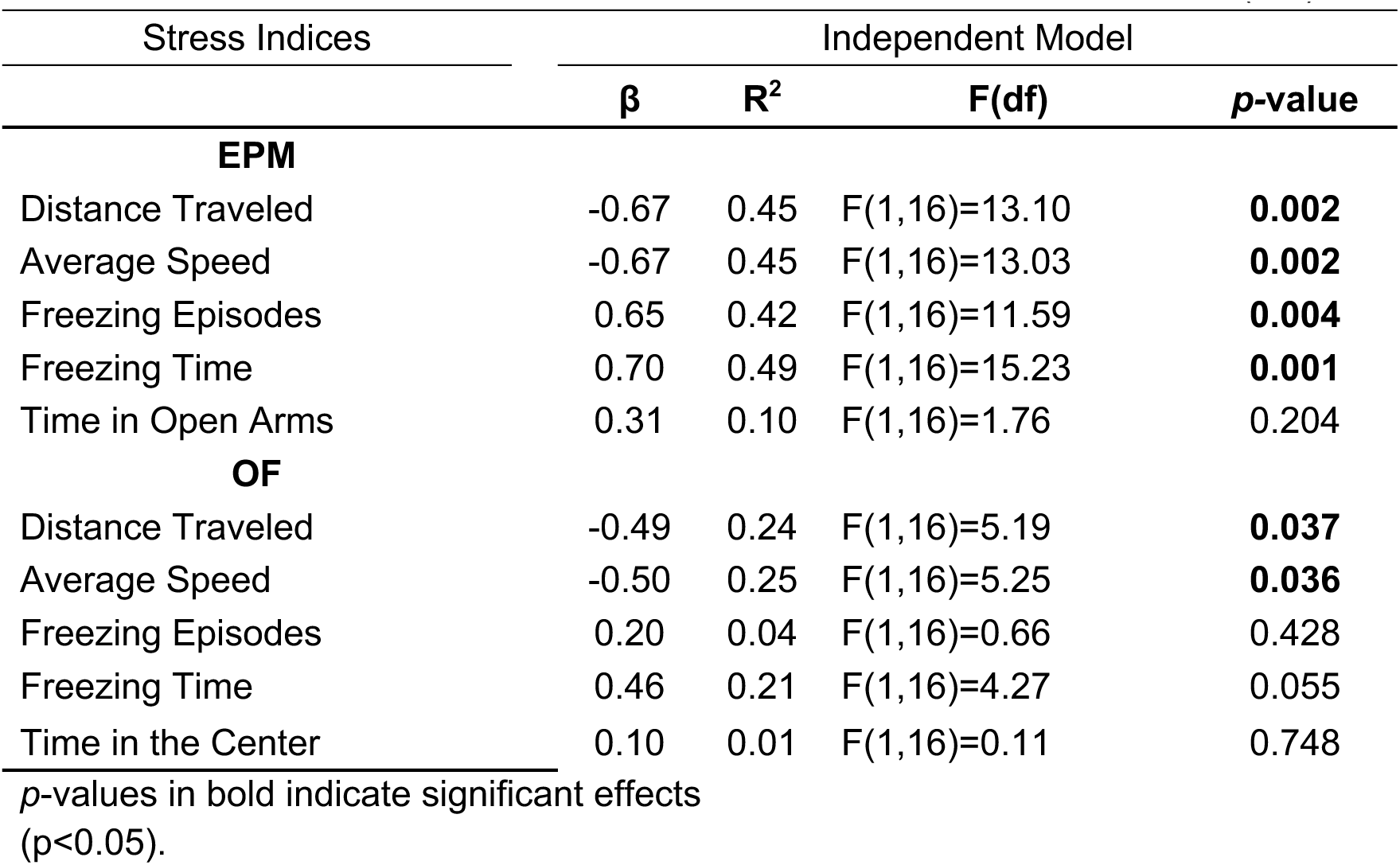
Association between stress indices and demand for ethanol (*EV*).

### 4.6. Concurrent ethanol and nicotine self-administration

An omnibus assessment of responding on active manipulanda for a reinforcer showed a significant effect of Condition [all conditions outlined in Table 1; (χ^2^_(3)_ = 8.87, 𝑝 = .031], a significant effect of a substance (nicotine vs ethanol; (χ^2^_(1)_ = 99.65, 𝑝 < .0001), and their interaction (χ^2^_(3)_ = 23.15, 𝑝 < .0001/. Given these findings, we proceeded to analyze responses for ethanol and nicotine in the presence of a secondary substance independently below.

#### The effect of nicotine on responding for ethanol

*<u>There</u>* was a significant effect of Condition on responding for ethanol +last 4 combinations in Table 1; (χ^2^_(3)_ = 23.79, 𝑝 < 0.0001/. Specifically, responding for ethanol was significantly increased when nicotine was concurrently available on FR1 schedule of reinforcement or when it was available noncontingently (Figure 5A).

**Fig. 5.**
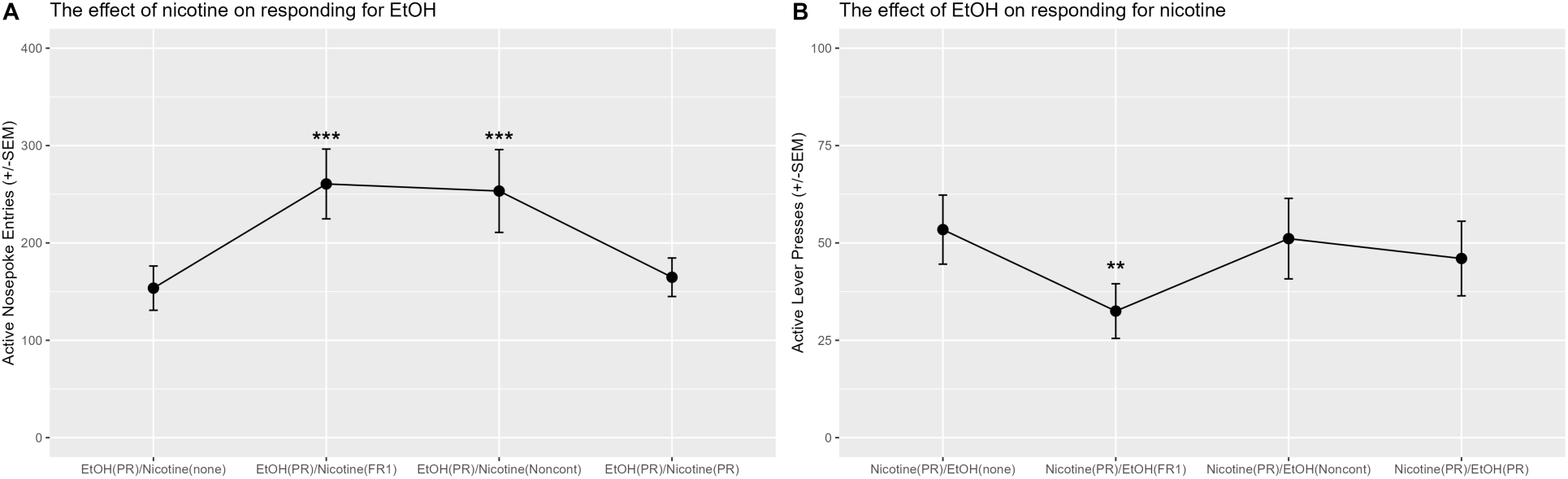
(A) The effect of nicotine on responding for ethanol on PR schedule of reinforcement. Nicotine increased responding for ethanol when nicotine was available on FR1 schedule of reinforcement or noncontingently. (B) The effect of ethanol on responding for nicotine on PR schedule of reinforcement. Ethanol decreased responding for nicotine when ethanol was available on FR1 schedule of reinforcement.

#### The effect of ethanol on responding for nicotine

There was a significant effect of Condition on responding for nicotine +first 4 combinations in Table 1; χ^2^_(3)_ = 12.40, p < 0.01/. Specifically, responding for nicotine was significantly lower when ethanol was concurrently available on the FR1 schedule of reinforcement in comparison to sessions when ethanol was not available [see the difference between Nicotine(PR)/EtOH(none) and Nicotine(PR)/EtOH(FR1) in Figure 5B].

### 4.7. Economic demand parameters predicting responding on PR schedule of reinforcement

Ethanol demand parameters predicted the response for ethanol on a progressive ratio (PR) schedule of reinforcement, with *EV*, *O_max_*, and *P_max_* each accounting for approximately 44-45 % of the variance in active lever presses for ethanol-alone (Table S1). Conversely, parameters from nicotine demand did not predict responses for ethanol-alone on a PR schedule (Table S2). In terms of nicotine demand, four out of five parameters accurately predicted responses for nicotine on a PR schedule. Each parameter accounted for 37-70 % of the variance individually, and together, they explained 81 % of the variance (refer to Table S3). However, ethanol demand parameters did not predict responses for nicotine on the PR schedule (Table S4).

Furthermore, when nicotine was available on a fixed ratio 1 (FR1) schedule, or administered non-contingently, its demand parameters did not predict responses for ethanol on a PR schedule (Tables S5 and S6, respectively). Similarly, ethanol demand parameters did not predict responses for nicotine on a PR schedule when ethanol was administered non-contingently (Table S7). These results underscore the utility of the PR schedule of reinforcement in assessing individual differences in effort allocation for the self-administration of substances like nicotine and ethanol. Moreover, these results contribute to our understanding of the interaction dynamics between commonly used substances in a closed economy setting, suggesting that individual-level assessment could be a useful strategy for evaluating complex interactions.

## 5. Discussion

Nicotine and ethanol co-abuse often leads to rapid dependency, adverse health impacts, and high rates of preventable mortality (Britt and Bonci, 2013, 2013; DiFranza and Guerrera, 1990; Littleton et al., 2007; Organization, 2017, 2017; Peacock et al., 2018). Previous studies show that nicotine and ethanol can influence each other’s rewarding and reinforcing effects, but systematic assessments mirroring clinical usage are lacking. Past preclinical research has enhanced our understanding of the combined neurobiological and behavioral effects of these substances, yet individual effects remain understudied. Addressing this gap, our study investigated nicotine and ethanol interactions using a long-access self-administration model, with a focus on both grouped and individual data. This comprehensive study initially assessed individual economic demand for sucrose, sweetened ethanol, ethanol alone, and nicotine, establishing a baseline for each rat. After the ethanol self-administration phase, withdrawal from ethanol was assessed. Subsequently, we explored the interactive effects of nicotine and ethanol, assessing how the availability and cost of one substance affected the consumption of the other under various reinforcement schedules. In our study, we found clear patterns and relationships both at the group and individual levels, which offer a better understanding of ethanol and nicotine use dynamics. Group-level observations revealed distinct economic demand patterns for sucrose, sweetened ethanol, and ethanol-alone, with sucrose evoking the highest demand and ethanol-alone showing increased sensitivity to cost changes under our experimental conditions. However, when viewed from an individual level, relationships between demand indices emerged, with demand for sucrose predicting that for sweetened ethanol and shared sensitivity to price increases observed between sweetened ethanol and ethanol alone.

Importantly, we also identified a similar individual-level sensitivity to cost changes between ethanol and nicotine, suggesting a potential link in their price response patterns. Another critical individual-level observation was the connection between stress indices and ethanol demand, where the economic demand for ethanol could forecast stress-related behavioral responses during withdrawal. Finally, at a group level, the concurrent self-administration of ethanol and nicotine showed reciprocal effects, with reduced responding for nicotine in the presence of ethanol and increased responding for ethanol in the presence of nicotine. Our supplemental analysis showed that the demand parameters could predict individual responses on a progressive ratio (PR) schedule of reinforcement for both ethanol and nicotine, highlighting the utility of PR schedules in assessing individual differences in effort allocation for self-administration. Thus, our study sheds light on the interactions between nicotine and ethanol, highlighting significant differences in demand for different substances, relationships between demand indices, and reciprocal effects during concurrent self-administration.

Although substantial efforts have been dedicated to preclinical research aimed at understanding the etiology of substance use disorder and developing effective treatment strategies, a notable gap remains in translating this research into cessation and relapse prevention treatments. The effectiveness of currently available treatments is challenging to determine due to variations in inclusion criteria and observation durations across clinical studies (Alpert et al., 2013; Bottlender and Soyka, 2005; Le Foll et al., 2014; Le Strat et al., 2011; Nunes et al., 2018). For instance, low motivation to quit or low consumption levels often lead to participant exclusions (Alpert et al., 2013; Le Strat et al., 2011). However, the effectiveness of these treatments is modest at best. One contributing factor to the limited translation from “bench to bedside” is the distinct approach to subject selection between clinical studies and grouped preclinical experimental designs. Clinical studies often recruit individuals with a significant substance use history and high motivation to quit, whereas preclinical studies typically utilize supposedly homogeneous samples (e.g., outbred rodents) randomly assigned to experimental conditions, treating within-group variance as an error. To enhance external validity, preclinical studies may consider investigating individual differences across various phases of the substance use continuum. Investigating individual effects can provide insights into prognostic and predictive markers associated with substance use disorder. This individual-centric method may hold the promise of more potent and personalized strategies, paving the way to more effective solutions for substance use disorder.

Our current study was designed to better understand individual differences in responding for ethanol and nicotine during the drug-taking phase. Specifically, we designed a study where rats first self-administer ethanol, and their economic demand for ethanol is assessed using a reinforcer demand modeling. Rats are then equipped with intravenous catheters, allowed to self-administer nicotine, and their economic demand for nicotine is also assessed using a reinforcer demand modeling. In the final phase of the study, rats are allowed to self-administer ethanol and nicotine concurrently, and the effect of one substance on the rate of responding for another substance is assessed using a PR schedule of reinforcement. Because catheter patency usually can be maintained only for a limited period of time (30 – 45 days) and because the acquisition of ethanol self-administration using a fading protocol usually takes much longer than the acquisition of nicotine self-administration, we elected to start with the assessment of economic demand for ethanol first and then to progress to the assessment of economic demand for nicotine and subsequently to a concurrent drug administration. Because we elected to use sucrose fading for the acquisition of ethanol self-administration, this allowed us also to acquire economic demand for sucrose alone and for sweetened ethanol. Having a record of individual economic demand for sucrose alone and for sweetened ethanol allows asking deeper questions about the relationships between economic demand for a food reinforcer like sucrose, sweetened ethanol, and ethanol alone. For example, we were able to show that the economic demand for sucrose can largely predict the economic demand for sweetened ethanol.

Specifically, our results indicate that rats that work hard for sucrose in our experimental conditions also work hard for sweetened ethanol. This finding suggests that the individual preference for food reinforcement can drive a preference for sweetened ethanol that models calorie-enriched alcoholic beverages like beer or mixed drinks. We also showed that the elasticity of demand for sweetened ethanol predicts responding for ethanol alone. This relationship suggests that individuals that show persisted responding for sweetened ethanol in the face of price increases are also insensitive to price change for ethanol alone. Importantly, because our design treats economic demand for a reinforcer as a continuous variable, we also show that rats that do not find sucrose highly reinforcing also do not find sweetened ethanol highly reinforcing and that rats that are sensitive to the price change for sweetened ethanol are also sensitive to the price change for ethanol alone. Lastly, the data from this phase of our study reveals that the economic demand for sucrose does not correlate with the demand for ethanol alone. This suggests that increased sensitivity to food-based reinforcement doesn’t necessarily extend to ethanol reinforcement within the constraints of our experimental setup.

Behavioral economics postulates that the amount of effort exerted for a reinforcer is dictated by its price. Simply put, the total consumption tends to decrease as the price of the reinforcer increases (Allison, 1979; Bickel et al., 1992; Hursh, 2014). Furthermore, the relationship between consumption and price is typically expressed by the elasticity of economic demand, which measures consumption rates across varying prices. For instance, if a reinforcer’s consumption diminishes with increasing prices, its demand is considered elastic. Conversely, if consumption remains stable despite a price hike, the demand is deemed inelastic. Essential items such as bread, milk, or gasoline usually exhibit inelastic demand because they are necessities. Consequently, consumers are likely to buy them despite significant price increases. Likewise, addictive substances are often viewed as having inelastic demand, as consumers regard them as essentials, being willing to pay a substantial amount to acquire them (Hursh, 1984; Hursh et al., 2005; Schwartz et al., 2021). In an open economy, characterized by a variety of goods available in different categories, the price of one commodity can influence the consumption of others. Within this system, some commodities can substitute for others, some may complement others, and some are independent. For example, goods are considered substitutes if consumers view them as similar or interchangeable, thus reducing the consumption of one if another is readily available. Conversely, a complementary good’s consumption increases when its counterpart’s consumption also increases, while an independent good’s consumption is unaffected by the consumption of other goods. Previous research examining the relationship between ethanol and other reinforcers suggested that ethanol and sucrose might function as substitutable reinforcers, as limiting the availability of sucrose increases ethanol consumption (Samson and Lindberg, 1984). However, later studies demonstrated that sweetened ethanol is more reinforcing than sucrose alone. This is because pre-session feeding decreases the response maintained by sucrose but not by ethanol (Heyman, 1993), the demand for sweetened ethanol was more inelastic compared to sucrose alone (Petry and Heyman, 1995), and increases in the price of sucrose or its alternatives systematically decreased consumption, whereas similar price increases for sweetened ethanol did not reduce consumption similarly (Heyman, 2000, 1997; Heyman et al., 1999; Kim and Kearns, 2019; Petry and Heyman, 1995). Our study builds upon these prior findings by systematically comparing the economic demands for sucrose, sweetened ethanol, and ethanol alone. We incorporated a comprehensive approach that includes the primary associated demand indexes and levels of assessment at both grouped and individual levels. With this approach, we revealed that rats were more inclined to work harder for sucrose than for sweetened ethanol or ethanol alone (refer to *EV*; see Figures 2 and 3A). Additionally, the demand for sucrose was more inelastic compared to that for ethanol alone. Contrary to previous findings, our results, within the context of our experimental design, suggest that sucrose has a stronger reinforcing effect than sweetened ethanol, and rats display less sensitivity to price hikes for sucrose compared to ethanol alone. The discrepancy with prior studies likely arises from differences in experimental design, assessment methods, and variations in session duration between earlier studies and our current study (with short-access primarily used in early studies vs. long-access employed in our study).

We further expand on prior studies that compared the grouped reinforcing values of sucrose, sweetened ethanol, and ethanol alone by evaluating the individual effects linked to the economic demand for these reinforcers. For the first time, we demonstrate that an individual’s demand for sucrose can predict the economic demand for sweetened ethanol. Specifically, we illustrate that the *EV*, *α*, *O_max_*, and *P_max_* indices for sucrose can all predict a corresponding demand index for sweetened ethanol, indicating a positive correlation between these measures. Consequently, our findings suggest that rats demonstrating a high sensitivity to sucrose reinforcement (indicated by a higher *EV*) are also highly sensitive to sweetened ethanol reinforcement. To operationalize this, rats working harder for sucrose also work hard for sweetened ethanol, suggesting that in these individuals, consumption of sweetened ethanol is partially driven by the reinforcing attributes of sucrose. On the other hand, our findings indicate that rats that exert more effort for sweetened ethanol do not necessarily show a high preference for ethanol alone, as the economic demand for sweetened ethanol does not directly correlate with responses for pure ethanol (refer to Table 2). It’s crucial to note that these effects emerge within the context of the long-access self-administration model for all these substances. Our data shows that within our experimental design, ethanol self-administration induces repeated withdrawal effects, as evidenced by performance in behavioral tests associated with stress and anxiety (refer to Figure 4). Furthermore, our data reveal a significant association between the economic demand for ethanol and performance on seven out of ten metrics gathered from stress and anxiety-related tests, suggesting that rats with a higher demand for ethanol experience a more pronounced magnitude of withdrawal effects throughout the study (refer to Table 3). Overall, our findings show that at the individual level, rats that find sucrose highly reinforcing also find sweetened ethanol highly reinforcing, and this subset of rats may be different from those who find pure ethanol highly reinforcing. Moreover, it is possible that ethanol withdrawal effects contribute to how hard some rats are willing to work for ethanol and thus constitute another dimension of reinforcing effects associated with ethanol reward.

Previous studies have employed a variety of experimental approaches to examine the relationship between ethanol and nicotine use. The disparities in administration models (contingent vs. noncontingent), routes of administration (drinking solutions vs. vapor vs. systemic injections vs. intravenous self-administration), and session length (short-access vs. long-access) likely contribute to the variability in reported effects. With this in mind, our current understanding of the interaction effects between ethanol and nicotine in the preclinical field remains limited. This motivated us to design a study that would begin to evaluate this interaction using a preclinical model relevant to the patterns of substance use observed in humans. Our study unveiled some novel findings that enhance our understanding of the relationship between ethanol and nicotine use. We identified significant group-level differences in the elasticity of economic demand for ethanol and nicotine. Specifically, the demand for ethanol was more elastic than that for nicotine (see Figure 3B). This indicates that responses for ethanol-alone were more sensitive to cost changes compared to those for nicotine-alone. Exploring individual elasticity, we found that rats sensitive to cost changes for ethanol displayed a similar sensitivity to cost changes for nicotine (see *α* section in Table 2). This also suggests that rats resilient to cost changes for ethanol demonstrated similar resilience to cost changes for nicotine. These results not only underscore a potential connection in price response patterns between ethanol and nicotine, but they also highlight the importance of considering individual differences when evaluating substance use behavior. Our interaction tests showed that both contingent “low-cost” and noncontingent nicotine administration increased responses for ethanol (see Figure 5A). It is crucial to note that there was a significant difference in the effect of contingent “low-cost” and “high-cost” nicotine availability on the response for ethanol. This is because the consumption of ethanol increased only when nicotine was available on the FR1 schedule of reinforcement (see Figure 5A). Our interaction tests also revealed that contingent “low-cost” ethanol significantly decreased responding for nicotine on a progressive schedule of reinforcement, while noncontingent nicotine or contingent “high-cost” nicotine availability had no effect (Figure 5B).

To our knowledge, this is the first demonstration of the effects of ethanol on responding for nicotine using a model where both substances are concurrently available for self-administration during the session. Altogether, our study illustrates that it is feasible to study the interaction effects between ethanol and nicotine using a model where both substances are self-administered separately and in a manner that mirrors human use (e.g., routes of administration and increased daily access). Overall, our findings suggest that the contingency and “cost” of a co-administered substance may significantly influence the interaction effects associated with polydrug use, or at the very least, the interaction effects between ethanol and nicotine. Additional studies will be necessary to confirm our results and expand our findings to other commonly used substances.

Tobacco and alcohol are significant contributors to preventable mortality worldwide. Co-use of these substances has been associated with accelerated dependence development, a broader range of negative health outcomes, and increased difficulty in quitting (Devlin and Henry, 2008; Frie et al., 2021; Hurt et al., 1996; Kohut, 2017; McKee and Weinberger, 2013; Mello et al., 1980; Mintz et al., 1985). Therefore, it is important to understand how these substances interact with each other under various conditions. In our study, we aimed to establish a model that may help to investigate the co-use of ethanol and nicotine in the preclinical setting. We examined the interaction effects of ethanol and nicotine by employing a reinforcer demand modeling and a combination of self-administration tests, which involved progressive ratio schedules of reinforcement and noncontingent administration of complementary substances. By utilizing these approaches, we aimed to better understand the dynamics between ethanol and nicotine use in terms of their reinforcing properties and demand characteristics in specific contexts like the one we created in our study. Our approach allowed us to address various research questions in a single study and analyze data at both grouped and individual levels. Importantly, our study highlighted the utility of individual data in predicting behaviors across different phases of the substance use cycle. Although this type of study is resource and time-demanding, we believe it is important to continue investigating polydrug use with the help of comprehensive models that can provide a broad spectrum of grouped and individual data related to different facets of polydrug use. We also believe that these types of studies that focus on individual data may benefit from a much larger sample size than what is currently practiced in the preclinical field to assess grouped effects. Additional studies focusing on sex differences and individual effects associated with pharmacological interventions and on defining vulnerable endophenotypes are also needed to continue expanding our understanding of ethanol and nicotine use comorbidity. Finally, understanding how an individual’s reinforcement history and polydrug use interact is essential for creating effective, personalized treatments and improving treatment outcomes for those with co-existing ethanol and nicotine use disorders.

## Acknowledgements

This research and S. Charntikov were supported by NIGMS (GM113131).

## 7. Supplemental Tables

**Table S1.**
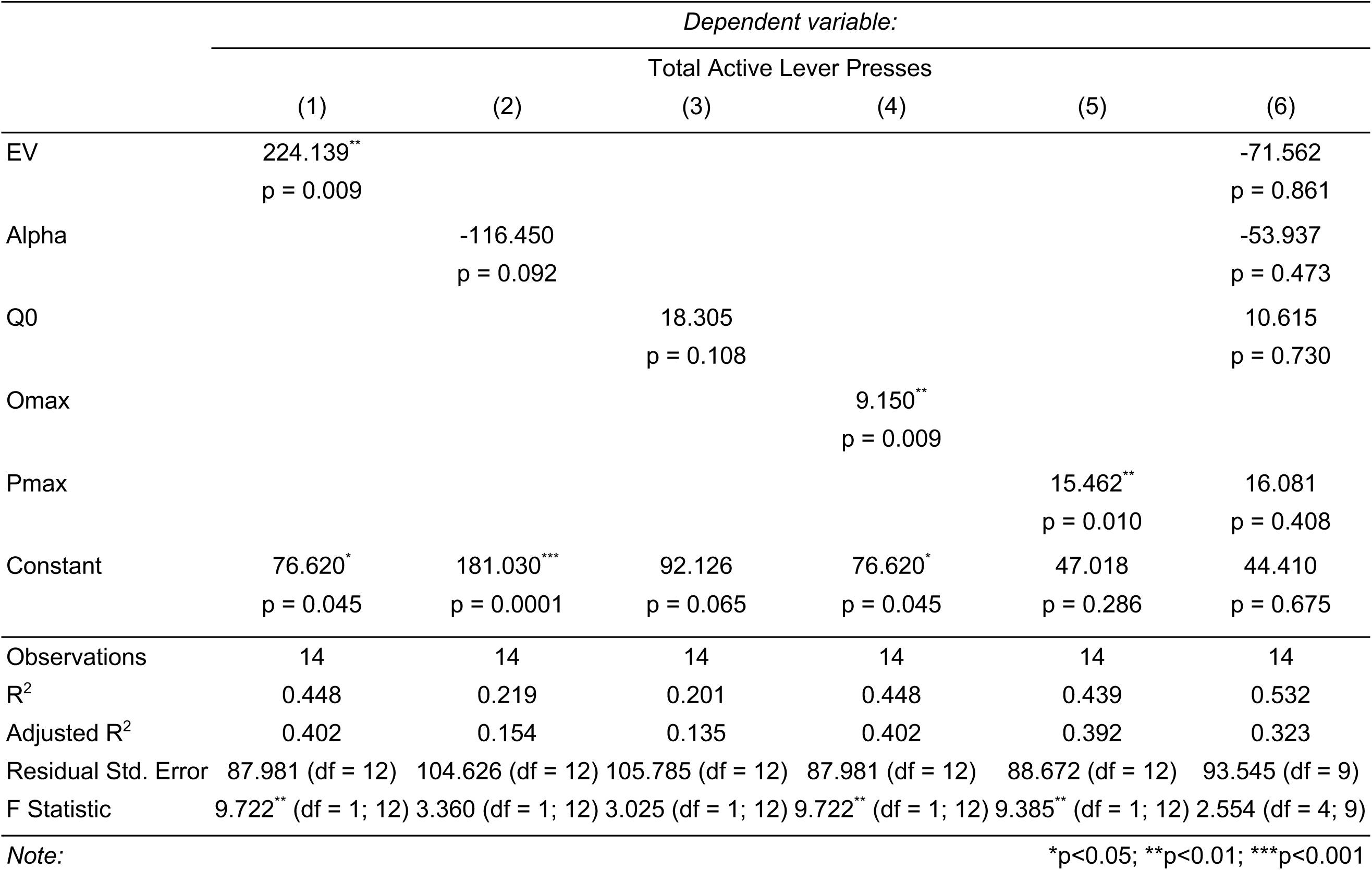
Ethanol demand parameters predict responding for ethanol on the PR schedule of reinforcement

**Table S2.** Nicotine demand parameters do not predict responding for ethanol on PR schedule of reinforcement

|  | *Dependent variable:* |  |  |  |  |  |
| --- | --- | --- | --- | --- | --- | --- |
|  | Total Active Lever Presses |  |  |  |  |  |
|  | (1) | (2) | (3) | (4) | (5) | (6) |
| EV | 52.178<br>p = 0.575 |  |  |  |  | 23.330<br>p = 0.926 |
| Alpha |  | 160.164<br>p = 0.915 |  |  |  | 1,249.787<br>p = 0.577 |
| Q0 |  |  | -0.251<br>p = 0.979 |  |  | 5.340<br>p = 0.667 |
| Omax |  |  |  | 1.877<br>p = 0.575 |  |  |
| Pmax |  |  |  |  | 0.968<br>p = 0.470 | 1.650<br>p = 0.633 |
| Constant | 131.796*<br>p = 0.017 | 150.533**<br>p = 0.002 | 153.899**<br>p = 0.004 | 131.796*<br>p = 0.017 | 131.871**<br>p = 0.008 | 70.519<br>p = 0.539 |
| Observations | 13 | 13 | 13 | 13 | 13 | 13 |
| R<sup>2</sup> | 0.030 | 0.001 | 0.0001 | 0.030 | 0.049 | 0.103 |
| Adjusted R<sup>2</sup> | -0.059 | -0.090 | -0.091 | -0.059 | -0.038 | -0.345 |
| Residual Std. Error | 102.521 (df = 11) | 104.014 (df = 11) | 104.068 (df = 11) | 102.521 (df = 11) | 101.511 (df = 11) | 115.561 (df = 8) |
| F Statistic | 0.335 (df = 1; 11) | 0.012 (df = 1; 11) | 0.001 (df = 1; 11) | 0.335 (df = 1; 11) | 0.562 (df = 1; 11) | 0.230 (df = 4; 8) |
| *Note:* |  |  |  |  | *p<0.05; **p<0.01; ***p<0.001 |  |

**Table S3.** Nicotine demand parameters predict responding for nicotine on the PR schedule of reinforcement

|  | *Dependent variable:* |  |  |  |  |  |
| --- | --- | --- | --- | --- | --- | --- |
|  | Total Nosepoke Entries |  |  |  |  |  |
|  | (1) | (2) | (3) | (4) | (5) | (6) |
| EV | 112.180***<br>p = 0.001 |  |  |  |  | 149.956<br>p = 0.139 |
| Alpha |  | -1,116.411<br>p = 0.060 |  |  |  | -190.594<br>p = 0.673 |
| Q0 |  |  | -7.465*<br>p = 0.036 |  |  | -4.807<br>p = 0.075 |
| Omax |  |  |  | 4.034***<br>p = 0.001 |  |  |
| Pmax |  |  |  |  | 1.392**<br>p = 0.003 | -0.909<br>p = 0.450 |
| Constant | 19.126<br>p = 0.105 | 78.187***<br>p = 0.0003 | 80.672***<br>p = 0.0002 | 19.126<br>p = 0.105 | 28.710*<br>p = 0.022 | 43.273<br>p = 0.085 |
| Observations | 12 | 12 | 12 | 12 | 12 | 12 |
| R<sup>2</sup> | 0.696 | 0.311 | 0.372 | 0.696 | 0.625 | 0.815 |
| Adjusted R<sup>2</sup> | 0.665 | 0.242 | 0.310 | 0.665 | 0.587 | 0.709 |
| Residual Std. Error | 24.206 (df = 10) | 36.434 (df = 10) | 34.769 (df = 10) | 24.206 (df = 10) | 26.881 (df = 10) | 22.569 (df = 7) |
| F Statistic | 22.872*** (df = 1; 10) | 4.510 (df = 1; 10) | 5.932* (df = 1; 10) | 22.872*** (df = 1; 10) | 16.654** (df = 1; 10) | 7.704* (df = 4; 7) |

Note: \*p<0.05; \*\*p<0.01; \*\*\*p<0.001

**Table S4.** Ethanol demand parameters do not predict responding for nicotine on the PR schedule of reinforcement

|  | *Dependent variable:* |  |  |  |  |  |
| --- | --- | --- | --- | --- | --- | --- |
|  | Total Nosepoke Entries |  |  |  |  |  |
|  | (1) | (2) | (3) | (4) | (5) | (6) |
| EV | -4.678<br>p = 0.930 |  |  |  |  | -339.004<br>p = 0.180 |
| Alpha |  | -12.065<br>p = 0.754 |  |  |  | 2.481<br>p = 0.961 |
| Q0 |  |  | 1.018<br>p = 0.827 |  |  | 28.615<br>p = 0.198 |
| Omax |  |  |  | -0.191<br>p = 0.930 |  |  |
| Pmax |  |  |  |  | 2.231<br>p = 0.633 | 5.972<br>p = 0.315 |
| Constant | 50.963*<br>p = 0.020 | 52.339**<br>p = 0.007 | 46.519*<br>p = 0.038 | 50.963*<br>p = 0.020 | 37.255<br>p = 0.221 | 8.654<br>p = 0.852 |
| Observations | 12 | 12 | 12 | 12 | 12 | 12 |
| R<sup>2</sup> | 0.001 | 0.010 | 0.005 | 0.001 | 0.024 | 0.263 |
| Adjusted R<sup>2</sup> | -0.099 | -0.089 | -0.094 | -0.099 | -0.074 | -0.158 |
| Residual Std. Error | 45.032 (df = 10) | 44.817 (df = 10) | 44.936 (df = 10) | 45.032 (df = 10) | 44.514 (df = 10) | 46.225 (df = 7) |
| F Statistic | 0.008 (df = 1; 10) | 0.104 (df = 1; 10) | 0.051 (df = 1; 10) | 0.008 (df = 1; 10) | 0.243 (df = 1; 10) | 0.625 (df = 4; 7) |

Note: \*p<0.05; \*\*p<0.01; \*\*\*p<0.001

**Table S5.** Nicotine demand parameters do not predict responding for ethanol on the PR schedule of reinforcement when nicotine is available on FR1

|  | *Dependent variable:* |  |  |  |  |  |
| --- | --- | --- | --- | --- | --- | --- |
|  | Total Active Lever Presses |  |  |  |  |  |
|  | (1) | (2) | (3) | (4) | (5) | (6) |
| EV | 187.411<br>p = 0.243 |  |  |  |  | -694.670<br>p = 0.240 |
| Alpha |  | -2,469.658<br>p = 0.295 |  |  |  | -2,986.862<br>p = 0.296 |
| Q0 |  |  | -26.007<br>p = 0.054 |  |  | -20.487<br>p = 0.193 |
| Omax |  |  |  | 6.740<br>p = 0.243 |  |  |
| Pmax |  |  |  |  | 3.123<br>p = 0.126 | 8.862<br>p = 0.233 |
| Constant | 215.944*<br>p = 0.015 | 329.111***<br>p = 0.0005 | 366.375***<br>p = 0.0001 | 215.944*<br>p = 0.015 | 214.602**<br>p = 0.006 | 458.746*<br>p = 0.017 |
| Observations | 11 | 11 | 11 | 11 | 11 | 11 |
| R<sup>2</sup> | 0.148 | 0.121 | 0.355 | 0.148 | 0.241 | 0.562 |
| Adjusted R<sup>2</sup> | 0.053 | 0.023 | 0.283 | 0.053 | 0.157 | 0.270 |
| Residual Std. Error | 151.409 (df = 9) | 153.786 (df = 9) | 131.719 (df = 9) | 151.409 (df = 9) | 142.891 (df = 9) | 132.958 (df = 6) |
| F Statistic | 1.563 (df = 1; 9) | 1.239 (df = 1; 9) | 4.957 (df = 1; 9) | 1.563 (df = 1; 9) | 2.860 (df = 1; 9) | 1.924 (df = 4; 6) |

*Note:* \*p<0.05; \*\*p<0.01; \*\*\*p<0.001

**Table S6.** Nicotine demand parameters do not predict responding for ethanol on the PR schedule of reinforcement when nicotine is administered noncontingently

|  | *Dependent variable:* |  |  |  |  |  |
| --- | --- | --- | --- | --- | --- | --- |
|  | Total Active Lever Presses |  |  |  |  |  |
|  | (1) | (2) | (3) | (4) | (5) | (6) |
| EV | 190.353<br>p = 0.283 |  |  |  |  | -656.208<br>p = 0.434 |
| Alpha |  | -1,238.463<br>p = 0.647 |  |  |  | -745.906<br>p = 0.851 |
| Q0 |  |  | -17.387<br>p = 0.282 |  |  | -6.868<br>p = 0.739 |
| Omax |  |  |  | 6.846<br>p = 0.283 |  |  |
| Pmax |  |  |  |  | 3.198<br>p = 0.157 | 10.629<br>p = 0.314 |
| Constant | 209.700*<br>p = 0.031 | 302.535**<br>p = 0.004 | 337.047**<br>p = 0.002 | 209.700*<br>p = 0.031 | 209.454*<br>p = 0.014 | 323.833<br>p = 0.182 |
| Observations | 10 | 10 | 10 | 10 | 10 | 10 |
| R<sup>2</sup> | 0.142 | 0.028 | 0.143 | 0.142 | 0.234 | 0.360 |
| Adjusted R<sup>2</sup> | 0.035 | -0.094 | 0.036 | 0.035 | 0.138 | -0.151 |
| Residual Std. Error | 166.815 (df = 8) | 177.629 (df = 8) | 166.763 (df = 8) | 166.815 (df = 8) | 157.637 (df = 8) | 182.227 (df = 5) |
| F Statistic | 1.327 (df = 1; 8) | 0.226 (df = 1; 8) | 1.333 (df = 1; 8) | 1.327 (df = 1; 8) | 2.445 (df = 1; 8) | 0.704 (df = 4; 5) |

*Note:* \*p<0.05; \*\*p<0.01; \*\*\*p<0.001

**Table S7.** Ethanol demand parameters do not predict responding for nicotine on the PR schedule of reinforcement when ethanol is available on FR1

|  | *Dependent variable:* |  |  |  |  |  |
| --- | --- | --- | --- | --- | --- | --- |
|  | Total Nosepoke Entries |  |  |  |  |  |
|  | (1) | (2) | (3) | (4) | (5) | (6) |
| EV | -35.236<br>p = 0.458 |  |  |  |  | 532.108<br>p = 0.323 |
| Alpha |  | -66.326<br>p = 0.445 |  |  |  | -510.050<br>p = 0.116 |
| Q0 |  |  | -1.438<br>p = 0.741 |  |  | -44.273<br>p = 0.297 |
| Omax |  |  |  | -1.438<br>p = 0.458 |  |  |
| Pmax |  |  |  |  | -4.374<br>p = 0.499 | -47.422<br>p = 0.183 |
| Constant | 46.886*<br>p = 0.039 | 39.649*<br>p = 0.015 | 41.259<br>p = 0.088 | 46.886*<br>p = 0.039 | 60.005<br>p = 0.144 | 345.824<br>p = 0.128 |
| Observations | 9 | 9 | 9 | 9 | 9 | 9 |
| R<sup>2</sup> | 0.081 | 0.086 | 0.017 | 0.081 | 0.068 | 0.683 |
| Adjusted R<sup>2</sup> | -0.050 | -0.045 | -0.124 | -0.050 | -0.065 | 0.365 |
| Residual Std. Error | 32.926 (df = 7) | 32.840 (df = 7) | 34.061 (df = 7) | 32.926 (df = 7) | 33.163 (df = 7) | 25.594 (df = 4) |
| F Statistic | 0.617 (df = 1; 7) | 0.657 (df = 1; 7) | 0.118 (df = 1; 7) | 0.617 (df = 1; 7) | 0.509 (df = 1; 7) | 2.152 (df = 4; 4) |

*Note:* \*p<0.05; \*\*p<0.01; \*\*\*p<0.001

## References

1. Acheson, A., Mahler, S.V., Chi, H., de Wit, H., 2006. Differential effects of nicotine on alcohol consumption in men and women. Psychopharmacology 186, 54. https://doi.org/10.1007/s00213-006-0338-y

2. Allison, J., 1979. Demand economics and experimental psychology. Behavioral Science 24, 403–415. https://doi.org/10.1002/bs.3830240606

3. Barrett, S.P., Tichauer, M., Leyton, M., Pihl, R.O., 2006. Nicotine increases alcohol self-administration in non-dependent male smokers. Drug and Alcohol Dependence 81, 197–204. https://doi.org/10.1016/j.drugalcdep.2005.06.009

4. Barrett, S.T., Thompson, B.M., Emory, J.R., Larsen, C.E., Pittenger, S.T., Harris, E.N., Bevins, R.A., 2020. Sex Differences in the Reward-Enhancing Effects of Nicotine on Ethanol Reinforcement: A Reinforcer Demand Analysis. Nicotine Tob Res 22, 238–247. https://doi.org/10/gnw7g8

5. Bickel, W.K., Hughes, J.R., De Grandpre, R.J., Higgins, S.T., Rizzuto, P., 1992. Behavioral economics of drug self-administration. Psychopharmacology 107, 211–216. https://doi.org/10.1007/BF02245139

6. Britt, J.P., Bonci, A., 2013. Alcohol and Tobacco: How Smoking May Promote Excessive Drinking. Neuron 79, 406–407. https://doi.org/10/gnf4ds

7. Burling, T.A., Ziff, D.C., 1988. Tobacco smoking: A comparison between alcohol and drug abuse inpatients. Addictive Behaviors 13, 185–190. https://doi.org/10/dvrd8d

8. Chandler, C.M., Maggio, S.E., Peng, H., Nixon, K., Bardo, M.T., 2020. Effects of ethanol, naltrexone, nicotine and varenicline in an ethanol and nicotine co-use model in Sprague-Dawley rats. Drug and Alcohol Dependence 212, 107988. https://doi.org/10/gnxdkw

9. Charntikov, S., Pittenger, S.T., Swalve, N., Barrett, S.T., Bevins, R.A., 2021. Conditioned enhancement of the nicotine reinforcer. Experimental and Clinical Psychopharmacology 29, 385–394. https://doi.org/10/gnw7bd

10. Deehan, G.A., Hauser, S.R., Waeiss, R.A., Knight, C.P., Toalston, J.E., Truitt, W.A., McBride, W.J., Rodd, Z.A., 2015. Co-Administration of Ethanol and Nicotine: The Enduring Alterations in the Rewarding Properties of Nicotine and Glutamate Activity within the Mesocorticolimbic System of Female Alcohol-Preferring (P) Rats. Psychopharmacology (Berl) 232, 4293–4302. https://doi.org/10/f7vzg4

11. Dermody, S.S., Tidey, J.W., Denlinger, R.L., Pacek, L.R., al’Absi, M., Drobes, D.J., Hatsukami, D.K., Vandrey, R., Donny, E.C., 2016. The impact of smoking very low nicotine content cigarettes on alcohol use. Alcoholism, Clinical and Experimental Research 40, 606–615. https://doi.org/10/gnw7bf

12. Devlin, R.J., Henry, J.A., 2008. Clinical review: Major consequences of illicit drug consumption. Crit Care 12, 1–7. https://doi.org/10.1186/cc6166

13. DiFranza, J.R., Guerrera, M.P., 1990. Alcoholism and smoking. J. Stud. Alcohol 51, 130–135. https://doi.org/10.15288/jsa.1990.51.130

14. Doyon, W.M., Dong, Y., Ostroumov, A., Thomas, A.M., Zhang, T.A., Dani, J.A., 2013. Nicotine Decreases Ethanol-Induced Dopamine Signaling and Increases Self-Administration via Stress Hormones. Neuron 79, 530–540. https://doi.org/10/f46kbz

15. Drugan, R.C., Eren, S., Hazi, A., Silva, J., Christianson, J.P., Kent, S., 2005. Impact of water temperature and stressor controllability on swim stress-induced changes in body temperature, serum corticosterone, and immobility in rats. Pharmacology, Biochemistry, and Behavior 82, 397–403. https://doi.org/10/bztpv3

16. European Monitoring Centre for Drugs and Drug Addiction, 2009. Polydrug use : patterns and responses. Publications Office, LU.

17. Frie, J.A., Nolan, C.J., Murray, J.E., Khokhar, J.Y., 2021. Addiction-Related Outcomes of Nicotine and Alcohol Co-use: New Insights Following the Rise in Vaping. Nicotine & Tobacco Research ntab231. https://doi.org/10/gnkvjk

18. Gilroy, S.P., Kaplan, B.A., Reed, D.D., 2020. Interpretation(s) of elasticity in operant demand. Journal of the experimental analysis of behavior 114, 106–115. https://doi.org/10/gnw677

19. Henningfield, J.E., Goldberg, S.R., 1983. Nicotine as a reinforcer in human subjects and laboratory animals. Pharmacology, Biochemistry, and Behavior 19, 989–992.

20. Heyman, G.M., 2000. An Economic Approach to Animal Models of Alcoholism. Alcohol Res Health 24, 132–139.

21. Heyman, G.M., 1997. Preference for saccharin-sweetened alcohol relative to isocaloric sucrose. Psychopharmacology 129, 72–78. https://doi.org/10.1007/s002130050164

22. Heyman, G.M., 1993. Ethanol regulated preference in rats. Psychopharmacology 112, 259–269. https://doi.org/10.1007/BF02244920

23. Heyman, G.M., Gendel, K., Goodman, J., 1999. Inelastic demand for alcohol in rats. Psychopharmacology 144, 213–219. https://doi.org/10.1007/s002130050996

24. Hursh, S.R., 2014. Behavioral economics and the analysis of consumption and choice, in: McSweeney, F.K., Murphy, E.S. (Eds.), The Wiley Blackwell Handbook of Operant and Classical Conditioning. John Wiley \& Sons, Ltd, Oxford, {UK}, pp. 275–305. https://doi.org/10.1002/9781118468135.ch12

25. Hursh, S.R., 1984. Behavioral Economics. Journal of the Experimental Analysis of Behavior 42, 435–452. https://doi.org/10.1901/jeab.1984.42-435

26. Hursh, S.R., Galuska, C.M., Winger, G., Woods, J.H., 2005. The economics of drug abuse: a quantitative assessment of drug demand. Molecular Interventions 5, 20–28. https://doi.org/10/c4jp9w

27. Hursh, S.R., Roma, P.G., 2016. Behavioral Economics and the Analysis of Consumption and Choice. Managerial and Decision Economics 37, 224–238. https://doi.org/10/gfxn2k

28. Hurt, R.D., Offord, K.P., Croghan, I.T., Gomez-Dahl, L., Kottke, T.E., Morse, R.M., Melton, L.J., III, 1996. Mortality Following Inpatient Addictions Treatment: Role of Tobacco Use in a Community-Based Cohort. JAMA 275, 1097–1103. https://doi.org/10.1001/jama.1996.03530380039029

29. Kazan, T., Charntikov, S., 2019. Individual differences in responding to bupropion or varenicline in a preclinical model of nicotine self-administration vary according to individual demand for nicotine. Neuropharmacology 148, 139–150. https://doi.org/10.1016/j.neuropharm.2018.12.031

30. Kazan, T., Robison, C.L., Cova, N., Madore, V.M., Charntikov, S., 2020. Assessment of individual differences in response to acute bupropion or varenicline treatment using a long-access nicotine self-administration model and behavioral economics in female rats. Behavioural Brain Research 385, 112558. https://doi.org/10/gnw7ft

31. Killeen, Peter R., and Kenneth W. Jacobs. “Coal Is Not Black, Snow Is Not White, Food Is Not a Reinforcer: The Roles of Affordances and Dispositions in the Analysis of Behavior.” The Behavior Analyst 40, no. 1 (June 1, 2017): 17–38. https://doi.org/10.1007/s40614-016-0080-7.

32. Kim, J.S., Kearns, D.N., 2019. Reduced ethanol self-administration in rats produced by the introduction of a high value non-drug alternative reinforcer. Pharmacology Biochemistry and Behavior 184, 172744. https://doi.org/10.1016/j.pbb.2019.172744

33. King, A., McNamara, P., Conrad, M., Cao, D., 2009. Alcohol-induced increases in smoking behavior for nicotinized and denicotinized cigarettes in men and women. Psychopharmacology 207, 107. https://doi.org/10.1007/s00213-009-1638-9

34. Kohut, S.J., 2017. Interactions between nicotine and drugs of abuse: A review of preclinical findings. Am J Drug Alcohol Abuse 43, 155–170. https://doi.org/10/gnxdsx

35. Korkosz, A., Taracha, E., Plaznik, A., Wrobel, E., Kostowski, W., Bienkowski, P., 2005. Extended blockade of the discriminative stimulus effects of nicotine with low doses of ethanol. European Journal of Pharmacology 512, 165–172. https://doi.org/10.1016/j.ejphar.2005.02.026

36. Kozlowski, L.T., Skinner, W., Kent, C., Pope, M.A., 1989. Prospects for smoking treatment in individuals seeking treatment for alcohol and other drug problems. Addictive Behaviors 14, 273–278. https://doi.org/10.1016/0306-4603(89)90058-0

37. Lallemand, F., Ward, R.J., De Witte, P., 2007. Nicotine increases ethanol preference but decreases locomotor activity during the initial stages of chronic ethanol withdrawal. Alcohol and Alcoholism 42, 207–218. https://doi.org/10.1093/alcalc/agm023

38. Lárraga, A., Belluzzi, J.D., Leslie, F.M., 2017. Nicotine Increases Alcohol Intake in Adolescent Male Rats. Frontiers in Behavioral Neuroscience 11, 25. https://doi.org/10/gnxdph

39. Le Foll, B., Goldberg, S.R., 2005. Ethanol does not affect discriminative-stimulus effects of nicotine in rats. European Journal of Pharmacology 519, 96–102. https://doi.org/10.1016/j.ejphar.2005.06.051

40. Littleton, J., Barron, S., Prendergast, M., Nixon, S.J., 2007. Smoking kills (alcoholics)! shouldn’t we do something about it? Alcohol and Alcoholism 42, 167–173. https://doi.org/10.1093/alcalc/agm019

41. Marshall, C.E., Dadmarz, M., Hofford, J.M., Gottheil, E., Vogel, W.H., 2003. Self-administration of both ethanol and nicotine in rats. Pharmacology 67, 143–149. https://doi.org/10.1159/000067801

42. McCarter, K.D., Li, C., Jiang, Z., Lu, W., Smith, H.A., Xu, G., Mayhan, W.G., Sun, H., 2017. Effect of Low-Dose Alcohol Consumption on Inflammation Following Transient Focal Cerebral Ischemia in Rats. Sci Rep 7, 12547. https://doi.org/10/gb2kr8

43. McKee, S.A., Weinberger, A.H., 2013. HOW CAN WE USE OUR KNOWLEDGE OF ALCOHOL-TOBACCO INTERACTIONS TO REDUCE ALCOHOL USE? Annu Rev Clin Psychol 9, 649–674. https://doi.org/10.1146/annurev-clinpsy-050212-185549

44. Mello, N.K., Mendelson, J.H., Sellers, M.L., Kuehnle, J.C., 1980. Effect of alcohol and marihuana on tobacco smoking. Clinical Pharmacology & Therapeutics 27, 202–209. https://doi.org/10.1038/clpt.1980.32

45. Mintz, J., Boyd, G., Rose, J.E., Charuvastra, V.C., Jarvik, M.E., 1985. Alcohol increases cigarette smoking: A laboratory demonstration. Addictive Behaviors 10, 203–207. https://doi.org/10.1016/0306-4603(85)90001-2

46. Mitchell, S.H., de Wit, H., Zacny, J.P., 1995. Effects of varying ethanol dose on cigarette consumption in healthy normal volunteers. Behavioural Pharmacology 6, 359–365. https://doi.org/10.1097/00008877-199506000-00006

47. Murray, C.J.L., Aravkin, A.Y., Zheng, P., Abbafati, C., Abbas, K.M., Abbasi-Kangevari, M., Abd-Allah, F., Abdelalim, A., Abdollahi, M., Abdollahpour, I., Abegaz, K.H., Abolhassani, 2020. Global burden of 87 risk factors in 204 countries and territories, 1990–2019: a systematic analysis for the Global Burden of Disease Study 2019. The Lancet 396, 1223–1249. https://doi.org/10.1016/S0140-6736(20)30752-2

48. Peacock, A., Leung, J., Larney, S., Colledge, S., Hickman, M., Rehm, J., Giovino, G.A., West, R., Hall, W., Griffiths, P., Ali, R., Gowing, L., Marsden, J., Ferrari, A.J., Grebely, J., Farrell, M., Degenhardt, L., 2018. Global statistics on alcohol, tobacco and illicit drug use: 2017 status report. Addiction 113, 1905–1926. https://doi.org/10/gdf7t8

49. Perkins, K.A., Fonte, C., Grobe, J.E., 2000. Sex differences in the acute effects of cigarette smoking on the reinforcing value of alcohol. Behavioural Pharmacology 11, 63–70. https://doi.org/10.1097/00008877-200002000-00007

50. Petry, N.M., Heyman, G.M., 1995. Behavioral Economics of Concurrent Ethanol-Sucrose and Sucrose Reinforcement in the Rat: Effects of Altering Variable-Ratio Requirements. Journal of the Experimental Analysis of Behavior 64, 331–359. https://doi.org/10.1901/jeab.1995.64-331

51. Pinheiro, J., Bates, D., DebRoy, S., Sarkar, D., (R CoreTeam), 2017. {nlme}: Linear and nonlinear mixed effects models.

52. Potthoff, A., Ellison, G., Nelson, L., 1983. Ethanol intake increases during continuous administration of amphetamine and nicotine, but not several other drugs. Pharmacology Biochemistry and Behavior 18, 489–493. https://doi.org/10/bqnd5x

53. R Core Team, 2019. R: A language and environment for statistical computing.

54. Samson, H.H., Lindberg, K., 1984. Comparison of sucrose-sucrose to sucrose-ethanol concurrent responding in the rat: Reinforcement schedule and fluid concentration effects. Pharmacology Biochemistry and Behavior 20, 973–977. https://doi.org/10.1016/0091-3057(84)90025-X

55. Schwartz, L.P., Blank, L., Hursh, S.R., 2021. Behavioral economic demand in opioid treatment: Predictive validity of hypothetical purchase tasks for heroin, cocaine, and benzodiazepines. Drug and Alcohol Dependence 221, 108562. https://doi.org/10/gnw682

56. Smith, B.R., Horan, J.T., Gaskin, S., Amit, Z., 1999. Exposure to nicotine enhances acquisition of ethanol drinking by laboratory rats in a limited access paradigm. Psychopharmacology 142, 408–412. https://doi.org/10/bpmcbk

57. Stafford, N.P., Kazan, T.N., Donovan, C.M., Hart, E.E., Drugan, R.C., Charntikov, S., 2019. Individual Vulnerability to Stress Is Associated With Increased Demand for Intravenous Heroin Self-administration in Rats. Frontiers in Behavioral Neuroscience 13, 134. https://doi.org/10/gnw7c2

58. Stuyt, E.B., 1997. Recovery Rates After Treatment for Alcohol/Drug Dependence: Tobacco Users vs. Non-Tobacco Users. American Journal on Addictions 6, 159–167. https://doi.org/10.3109/10550499709137027

59. Toneatto, A., Sobell, L.C., Sobell, M.B., Kozlowski, L.T., 1995. Effect of cigarette smoking on alcohol treatment outcome. Journal of Substance Abuse 7, 245–252. https://doi.org/10.1016/0899-3289(95)90008-X

60. Troisi, J.R., Dooley, T.F., Craig, E.M., 2013. The discriminative stimulus effects of a nicotine-ethanol compound in rats: Extinction with the parts differs from the whole. Behavioral Neuroscience 127, 899–912. https://doi.org/10.1037/a0034824

61. Venables, W.N., Ripley, B.D., Venables, W.N., 2002. Modern applied statistics with S. World Health Organization, 2018. Global status report on alcohol and health 2018. Geneva: World Health Organization.

62. World Health Organization, 2017. Who Report On The Global Tobacco Epidemic, 2017. World Health Organization.

